# CONSULT-II: Accurate taxonomic identification and profiling using locality-sensitive hashing

**DOI:** 10.1101/2023.11.07.566115

**Authors:** Ali Osman Berk Şapcı, Eleonora Rachtman, Siavash Mirarab

## Abstract

Taxonomic classification of short reads and taxonomic profiling of metagenomic samples are well-studied yet challenging problems. The presence of species belonging to ranks without close representation in a reference dataset is particularly challenging. While k-mer-based methods have performed well in terms of running time and accuracy, they tend to have reduced accuracy for such novel species. Here, we show that using locality-sensitive hashing (LSH) can increase the sensitivity of the k-mer-based search. Our method, which combines LSH with several heuristics techniques including soft LCA labeling and voting is, more accurate than alternatives in both taxonomic classification of individual reads and abundance profiling.

## Introduction

Metagenomic sequencing of microbial communities produces short DNA reads from unknown microorganisms (Handelsman, 2004), leading to a need for taxonomic identification based on reference datasets. One approach is to taxonomically identify reads and summarize the results to obtain the taxonomic profile of a sample, showing the relative abundances of taxonomic groups. However, despite the availability of mature read classification and profiling tools, benchmarking has revealed major gaps in the accuracy of existing methods (McIntyre *et al*., 2017; Meyer *et al*., 2019; Sczyrba *et al*., 2017; Ye *et al*., 2019). Precise identification is often hampered by the novelty of queries versus the genome-wide reference datasets and ambiguous matches. In addition, searching against large numbers of genomes is computationally demanding.

Taxonomic identification methods employ various strategies, including *k*-mer matching (Ames *et al*., 2013; Ounit et al., 2015; Wood *et al*., 2019; Lau et al., 2019; Lu et al., 2017), read mapping (Zhu *et al*., 2022), marker-based alignment (Liu *et al*., 2011; Milanese *et al*., 2019; Segata et al., 2012; Sunagawa et al., 2013), and phylogenetic placement (Asnicar *et al*., 2020; Shah et al., 2021; Truong *et al*., 2015). Regardless, they all essentially search for matches between reads in the sample and a reference set. The challenge is that a significant portion of the earth’s microbial diversity lacks close representatives in reference datasets (Choi *et al*., 2017), especially in poorly known microbial habitats like seawater or soil (Pachiadaki *et al*., 2019). Thus, most methods use some strategy to seek inexact matches between the query and references and use the results for classification and profiling.

Classification methods often exhibit reduced accuracy for *novel* sequences, which lack representation in reference sets (Liang *et al*., 2020; von Meijenfeldt *et al*., 2019; Nasko *et al*., 2018; Pachiadaki *et al*., 2019). For instance, Rachtman *et al*. (2020) found a leading tool, Kraken-II (Wood *et al*., 2019), faced significant degradation in domain-level classification as the genomic distance to the closest reference increased beyond 10%. Analyses of reads from less commonly sampled environments often fail at classification, even at the phylum-level (e.g., Pachiadaki *et al*., 2019). To tackle these challenges, efforts to build more dense reference sets are ongoing (McDonald *et al*., 2023; Parks *et al*., 2020; Wu *et al*., 2020), but these databases remain incomplete compared to the estimated 10^12^ microbial species (Locey and Lennon, 2016). Additionally, computational challenges arise in searching against large reference sets. Thus, we need accurate and scalable methods of identifying novel sequences with respect to distant reference genomes.

As reference sets grow larger, *k*-mer-based methods become more attractive than alignment-based approaches and phylogenetic placement due to their favorable balance between scalability, ease of use, and high accuracy (Ye *et al*., 2019; McIntyre *et al*., 2017). However, *k*-mer-based methods can be sensitive to reference set completeness if they only allow exact matches. The *k*-mer-based methods that rely on the presence/absence of long *k*-mers can accommodate novel sequences by allowing inexact matches. Kraken-II achieves this by masking some positions in a *k*-mer (default: 7 out of 31). Rachtman et al. (2021) showed that novel reads (e.g., those with 10–15% distance to the closest match) can be identified with higher accuracy by making inexact matches a central feature of the search. The resulting method, CONSULT, uses locality-sensitive hashing (LSH) to partition *k*-mers in the reference set into fixed-size buckets such that for a given *k*-mer, the reference *k*-mers with distance up to a certain threshold are within pre-determined buckets with high probability. By allowing inexact *k*-mer matching, CONSULT increased sensitivity without compromising precision in the contamination removal application (domain-level classification). However, CONSULT did not perform taxonomic identification at lower levels.

This paper adopts CONSULT and its increased *k*-mer matching sensitivity to the taxonomic classification problem. CONSULT estimates the Hamming distance (HD) between the query *k*-mer and its closest reference *k*-mers, a feature that Kraken-II lacks. Using the distances is the essence of our proposed approach, which we call CONSULT-II. To enable taxonomic classification, we need to track the reference genome(s) associated with each reference *k*-mer, a feature that CONSULT lacks and can require unrealistically large memory if done naively. We propose a probabilistic method to retain a single taxonomic ID per *k*-mer, making it possible to fit the database in the memory of modern server nodes. The next challenge is producing a single assignment based on potentially conflicting signals of different *k*-mers; we address this need using a weighted voting scheme that accounts for distances. Finally, we use a two-level normalization scheme for producing abundance profiles of complex samples using the votes directly. We evaluate the resulting method, CONSULT-II, using a large reference dataset in simulation studies, and show improved accuracy.

## Algorithm

### Background: CONSULT

The core idea of CONSULT is to find low HD matches efficiently using the bit-sampling LSH method (Har-Peled *et al*., 2012). The use of LSH for finding similar DNA sequences is not new (Berlin *et al*., 2015; Buhler, 2001; Rasheed et al., 2013; Luo et al., 2019). For example, Brown and Truszkowski (2013) addressed the related problem of phylogenetic placement using LSH to limit parts of the reference tree searched. The main focus and novelty of CONSULT, compared to existing work, is being able to search against a large number of reference sequences.

CONSULT tackles the following problem: Are there any *k*-mers in a given set of reference *k*-mers with Hamming distances (HD) less than some threshold *p* to a query *k*-mer? By default, CONSULT uses *k*=32 and *p*=3 (these are adjustable). CONSULT can feasibly index a large set of reference *k*-mers (e.g., 2^33^).

CONSULT employs two main data structures to represent a set of reference *k*-mers: an array *𝒦* that encodes each 32-mer as a 64-bit number, and *l*-many (default: *l*=2) fixed-sized hash tables **H**^1^, …, **H**^*l*^ with 4-byte pointers to *𝒦*; (and extra *n* + 1 bits when |*𝒦*| *>* 2^32+*n*^). Each hash table is a simple 2^2*h*^ × *b* matrix (default: *h*=15 and *b*=7) where each row is indexed by a hash value and the columns store pointers to *k*-mers in *𝒦*. For each hash table **H**^*i*^, we select *h* random but fixed positions of a 32-mer as its hash index. Thus, *k*-mer hashes are computed by simply extracting the corresponding bits from the 64-bit encoding of the *k*-mer, which is specifically designed to make these extractions efficient. For each query *k*-mer, the *l* hashes are computed, pointers from all ≤ *b* × *l* entries in the **H**^1^, …, **H**^*r*^ are followed to corresponding encodings in *𝒦*, and the HD is explicitly computed for each such encoding. CONSULT returns a match if there exist *k*-mers with distance ≤ *p*. As such, it has no false positive matches but false negatives (not finding a match) are possible. Using LSH, CONSULT limits the number of HD computations to a constant.

In our bit sampling scheme (Har-Peled *et al*., 2012), two *k*-mers at HD=*d* have the same hash with probability (1 − ^*d*^*/*_*k*_)^*h*^. Hence, given two independent *k*-mers, the probability that *at least one* of the hash functions is the same for both *k*-mers is given by

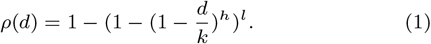

As desired, for *d* ≤ *p, ρ*(*d*) is close to 1. For *d* ≫ *p* and for some small enough *p* (e.g., *p*=3), it quickly drops to small values for several choices of *l* and *h*. Furthermore, since classification is done at the read level, we have *L* − *k* + 1 chances for a *k*-mer match (*L* = read length). While *k*-mer dependence across a read hampers computing the probability of having at least one LSH match between a read and a database (and independence assumption would be too inaccurate; see Fig. S1), we can still compute the expected number of such matches. Assuming the probability of a mismatch between each base pair of a read and a reference species is ^*d*^*/*_*k*_, the expected number of matching *k*-mers is (*L* − *k* + 1)*ρ*(*d*), which can be a large value for realistic settings (Fig. 1a). For example, with the default settings of *k*=32, *h*=15, *l*=2, for a 150bp read at 25% distance from the closest reference, we still expect 3.2 *k*-mer matches and can potentially classify it (assuming that *b* is large enough to fit all reference *k*-mers). Had we used *l* = 1 tables, this expected value would have been 1.6, making it likely to miss many such reads. If we assume 3–4 expected matches provide a sufficiently high probability of at least one match, *l* = 1 would suffice for *d* ≤ 0.21, while *l* = 3 and *l* = 4 would only increase our tolerance to *d* ≤ 0.27 and *d* ≤ 0.28. Given the linear increase in memory with *l*, we choose *l* = 2 as a tradeoff.

**Fig. 1:**
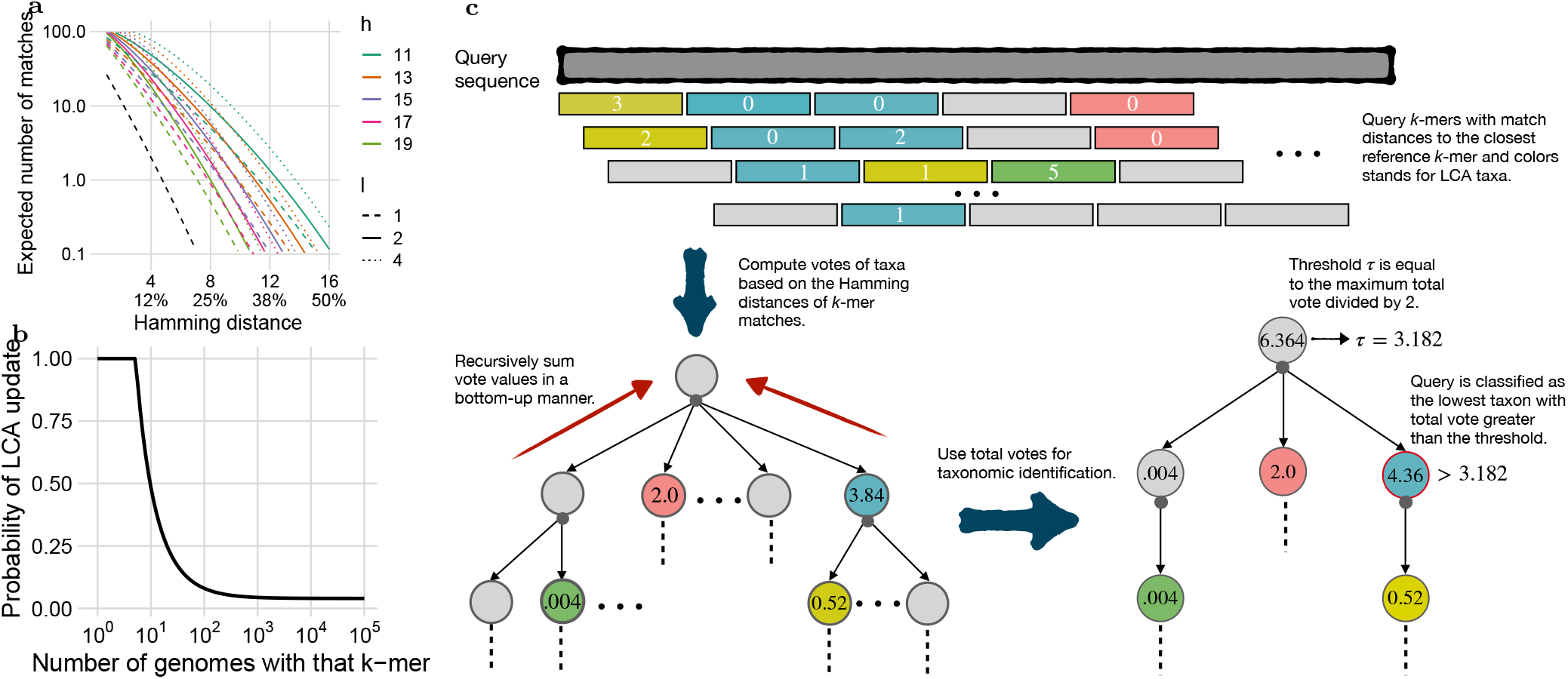
(a) The expected number of 32-mers matched by the LSH approach for a short read of length *L*=150 as the normalized distance ^*d*^*/*_*k*_ of the read to the closest match varies: (*L* − *k* + 1)*ρ*(*d*). Lines show different settings of *l* and *h* for an infinite *b*, i.e., all reference *k*-mers are stored in the library. The black line corresponds to *k*=35, *h*=35 − 7, and *l*=1, mimicking the default Kraken-II settings. (b) The probability of updating the LCA per each new *k*-mer goes down with the number of reference genomes in which that *k*-mer appears; we use *w*=4 and *s*=5 as shown. (c) Overview of the classification algorithm, consisting of three main stages: i) finding the closest inexact *k*-mer match using LSH ii) computing vote values based on the Hamming distances (HD) and aggregating vote values on the taxonomic tree iii) determining the most specific taxon, i.e., the lowest rank, above the threshold.

### Overview of the CONSULT-II changes versus CONSULT

To enable taxonomic classification, CONSULT needs to be extended to address several challenges. *i*) CONSULT was designed for a fixed reference library size. As a result, all the hashing settings (*h, l, b*) were fixed for a library of roughly 2^33^ 32-mers. To make the method more usable and flexible, it needs to be adjusted to the size of the input library. CONSULT-II uses several heuristics (see Şapcı *et al*. (2023) and supplementary Section S1) to estimate an efficient parameter configuration, as a function of the number of *k*-mers in the reference set and probability of matching two *k*-mers w.r.t. distance (*ρ*(*d*)). This heuristic enables adjusting the needed memory to reference size. *ii*) When building the reference library, we need to keep track of which *k*-mers belong to which set of taxonomic groups. Since keeping a fine-grained map will lead to an explosion of memory, we need heuristics (detailed below) to store *some* taxonomic information but also to keep the memory manageable. *iii*) At the query time, we need some way of combining all the inexact matches from all the *k*-mers of a read to derive a final identification and to summarize the results across all reads in a sample into a final profile.

### Soft lowest common ancestors (LCA) per *k*-mer

Considering the density of modern reference datasets, *k*-mers can appear frequently in several reference species, despite being long (e.g., *k* = 32). The required memory for keeping the associated set of taxonomic groups for each *k*-mer would quickly become infeasible. However, for ≤65536 taxonomic labels, keeping a single ID requires 2 bytes. Hence, storing one ID per *k*-mer would consume 16Gb for our standard libraries with 2^33^ *k*-mers, which is doable. We choose a single taxonomic ID representing a “soft” LCA of all the genomes that include the *k*-mer using the following procedure.

Let *N*_*i*_ denote the number of genomes that include *k*-mer *x*_*i*_ ∈ *𝒦*, which can be easily computed using pre-processing. At any point in time during the construction, a *k*-mer *x*_*i*_ ∈ *𝒦* is assigned a 2-byte taxonomic ID, denoted by *t*_*i*_. We process through each reference genome *g*; for each *k*-mer *x*_*i*_ of *g*, if it can be added to the database, we set or update the taxonomic ID *t*_*i*_ to be the LCA of the current *t*_*i*_ and species of *g* with probability

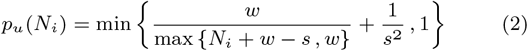

where *w* and *s* parameterize the rate of decrease and the offset of the probability function *p*_*u*_, respectively. We set *s*=5 and *w*=4 as the default values (see Fig. 1b). Note that the order of processing of the reference genomes has no significance, as every *k*-mer, including the first encountered, will be ignored with the same probability. Also, *k*-mers appearing in more than *s* genomes have a very small, but non-zero, probability of not having a taxonomic ID at all. The goal of the probabilistic soft LCA is to avoid pushing up taxonomic identifications due to errors in the reference libraries, as higher ranks are less informative. Imagine a *k*-mer that is found exclusively in 20 species of a particular genus, but is also found in one species of a completely different phylum. Using the hard LCA would push the *k*-mer up to the kingdom level, whereas the soft LCA will stay at the genus rank with 85% probability.

The probability function *p*_*u*_ is a heuristic without a theoretical ground but has two goals. First, it ensures *k*-mers are assigned an ID if they are *rare* among references (i.e., *N*_*i*_ ≤ *s*=5). Second, the probability of ignoring a genome smoothly increases as *N*_*i*_ grows. The ^1^*/s*^2^ term is to ensure that each *k*-mer has a non-zero probability of having a taxonomic ID associated with it, even if it is extremely common.

### Read level taxonomic identification

For each read, CONSULT-II produces a list of matched *k*-mers; and for each matched *k*-mer *x*, it outputs the soft LCA taxonomic ID and the distances between *x* and its closest match with the same hash index. To identify a read, we need to derive a single conclusion from all these potentially conflicting signals. We do so by considering each *k*-mer as providing a *vote* to the corresponding taxonomic ID, but weight votes by the match distance.

Let *𝒯* denote the set of all taxonomic IDs, and *𝒦* (*t*), *t* ∈ *𝒯* be all reference *k*-mers with *t* as their soft LCA. Each *k*-mer *x* in the set *ℛ* of query read *k*-mers might match multiple *k*-mers in the reference set *𝒦* with varying distances. A match of a lower distance should provide a strong signal. CONSULT-II accounts for this by giving a *k*-mer *x* ∈ *ℛ* a vote for the taxonomic ID *t* using:

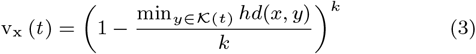

where *hd* gives the Hamming distance. The voting function (3) drops close to exponentially with distance min_*y*∈*𝒦* (*t*)_ *hd*(*x, y*). Computing (3) exactly is intractable due to the large size of *𝒦* (*t*). Instead, using LSH, we compare *x* only to *k*-mers *y* with the same hash index as *x*, finding matches with high probability. Moreover, we let *x* to vote for only a single taxonomic ID with the minimum distance (breaking ties arbitrarily). As LSH is not effective for high distances, we let a *k*-mer vote only if its minimum distance is below a threshold *d*_max_ (default: round(^3*p*^*/*_2_) =5). We set *d*_max_ to be higher than *p* because matches with distance above *p* might also be found; the LSH guarantees that *k*-mers with distance ≤ *p* are found with high probability, but more distant *k*-mers can also be found (see Fig. 1a).

Equation (3), however, is not enough because a vote for a child should also count towards parent ranks. We recursively sum up individual votes in a bottom-up manner using the taxonomic tree to derive a total vote value for each taxonomic ID:

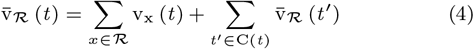

where C (*t* ) is the set of children of the taxon *t* .

By design, the votes 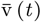 increase for higher ranks and reach their maximum at the root (Fig. 1c). To balance specificity and sensitivity, we require a majority vote. Let 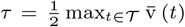. CONSULT-II classifies the read with the taxonomic ID 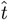 that belongs to the lowest rank satisfying the condition 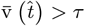. This choice of *τ* has a special property: Only a single taxonomic ID *t* can exceed *τ* at a given rank. Therefore, the taxonomic ID predicted by the described classification scheme is unique. Effectively, the classifier starts seeking a taxon at the lowest rank possible but also requires a certain level of confidence; hence, it immediately stops considering upper ranks once the vote value is large enough. Additionally, to avoid classification based on only high-distance matches, we require 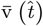 to be greater than some small threshold, which we explore in our experimental results.

### Taxonomic profiling

To derive taxonomic abundance profiles, instead of using read identifications, we use votes directly. For each taxonomic rank *r* (e.g., genus), we first normalize the total votes per read per rank, equalizing the contributions of each read to the profile (if it has any matches). For a read *ℛ* _*i*_, we simply set

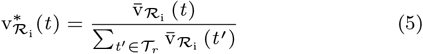

where *𝒯* _*r*_ is the set of all taxa at rank *r*. Next, we gather normalized total vote values of all *n* reads *ℛ*_1_, … *ℛ*n, in a sample, and normalize again to obtain the final profile. Let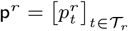 denote the relative abundance profile at rank *r*, summing up to 1. Then, we can set the relative abundance of taxon *t* to:

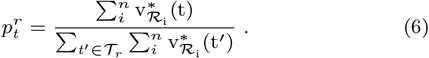

Here, 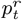 estimates the ratio of *reads* belonging to the taxon *t* in a given sample. Often, we are interested in the relative abundances of *cells* belonging to a taxon *t* (denoted by 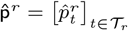, which needs incorporating genome sizes. We simply do so using:

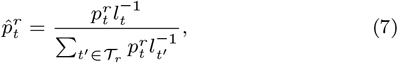

where *l*_*t*_ is the average genome length of all references in taxon *t*.

For both 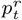 and 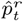, relative abundances sum up to 1. By default, CONSULT-II relaxes this constraint by including an *unclassified* taxon. This is achieved by propagating votes down to an artificial lineage that corresponds to the unclassified group, as each *k*-mer match to an LCA taxon provides evidence for its children – but it is unclear which. In other words, we augment the taxonomic tree by adding a lineage under each taxon, which continues until the species rank. Then, all votes to any non-species taxon are moved along this lineage to an artificial node at species rank. This is equivalent to changing the denominator of Eq. (5) with the total vote at the root of the taxonomic tree.

## Experimental setup

To benchmark CONSULT-II, we constructed reference libraries using the WoL microbial genomic dataset of Zhu *et al*. (2019), which is composed of 10,575 species and a reference phylogeny. Five genomes with IDs missing from NCBI were excluded. All methods were run with the same reference set. The hashing parameters of CONSULT-II were set to *h*=15, *b*=7, *l*=2, and *k*=32 (minimized from canonical 35-mers). For other parameters, default values were used: *w*=4, *s*=5 for LCA probability function *p*_*u*_ (Fig. 1b) and *d*_max_=5 for the vote function. We used default settings for Kraken-II, without masking low-complexity sequences, as Rachtman *et al*. (2020) found default settings to be preferable for query identification. We also constructed the CLARK database using the standard parameters, e.g., *k*=31, default classification mode, species rank for classification. Note that following Rachtman *et al*. (2021), 100 archaeal genomes were left out from the reference and used as *part of* the query set.

### Experiment 1: controlled novelty

We compared the classification performance of CONSULT-II with two popular methods: Kraken-II (Wood *et al*., 2019) and CLARK (Ounit *et al*., 2015), which are among the leading metagenomic identification tools based on benchmarking studies (Ye *et al*., 2019; Meyer *et al*., 2019; Sczyrba *et al*., 2017; McIntyre *et al*., 2017). Kraken-II maps each *k*-mer in a read to the LCA of all genomes that contain that *k*-mer and then counts the mapped *k*-mers on the taxonomic tree to infer a taxon prediction. CLARK is a supervised sequence classification method that again relies on exact *k*-mer matching. It uses the notion of discriminative *k*-mers to build a library of reference genomes. Here, we evaluate accuracy one read at a time, each simulated from a query genome.

Let the *novelty* of a query genome be defined as its minimum genomic nucleotide distance (i.e., one minus average nucleotide identity), as approximated by Mash (Ondov *et al*., 2016), to any genome in the reference database. We refer to this quantity as MinGND. We carefully selected query genomes to span a range of novelty (Fig. S2), expecting that more novel queries will be more challenging. We created two sets of queries: bacterial and archaeal. For the bacterial set, we selected 120 bacterial genomes among genomes added to RefSeq after WoL was constructed. Queries range from near-identical to reference genomes to very novel (e.g., 22 with MinGND *>* 0.22; Fig. S2). Query genomes span 29 phyla, and most queries are from distinct genera (102 genera across 120 queries); only two query genomes belong to the same species. The 100 archaeal queries were chosen by Rachtman *et al*. (2021) from WoL set using a similar approach and were excluded from the reference set. We generated 150bp synthetic reads using ART (Huang *et al*., 2012) at higher coverage, and then subsampled down to 66667 reads for each query (i.e., 10Mbp per sample).

We evaluated the predictions of each tool with respect to the NCBI taxonomy. For each read, we evaluate it separately at each taxonomic rank *r*. When the reference library had at least one genome matching the query taxon at rank *r*, we called it a *positive*: *T P* if a tool found the correct label, *FP* if it found an incorrect label, and *FN* if it did not classify at rank *r*. When the reference library did not have any genomes from the query taxon at rank *r*, we called it a *negative*: *T N* if a tool did not classify at rank *r, FP* if it classified it, which would necessarily be false. We show the precision ^*T P*^*/*(*T P* + *F P* ), the recall ^*T P*^*/*(*T P* + *F N* ), and F1 ^2*T P*^*/*(2*T P* + *F P* + *F N* ) which combines both sensitivity and specificity. We ignored queries at levels where the true taxonomic ID given by NCBI is 0, which indicates a missing rank.

### Experiment 2: abundance profiling

We also evaluated the ability of CONSULT-II to perform taxonomic abundance profiling using CAMI-I (Sczyrba *et al*., 2017) and CAMI-II (Meyer *et al*., 2022) benchmarking challenges. We compared tools using metrics provided by the open-community profiling assessment tool (OPAL) (Meyer *et al*., 2019). For CONSULT-II, we allowed unclassified taxa in the profile.

CAMI-I dataset contains five different high-complexity samples, each of size 75Gbp, which are simulated to mimic the abundance distribution of the underlying microbial communities. Among many metrics, we chose two metrics singled out in the original OPAL paper: the Bray-Curtis dissimilarity between the estimated profile and the true profile and Shannon’s equitability as a measure of alpha diversity. We report the summary of these two metrics across five samples. We use CAMI-I dataset for empirical evaluation of our method’s heuristics. Here, we used the same reference libraries constructed for controlled novelty experiments from the WoL dataset. As a result, we include only Bracken (Lu *et al*., 2017) and CLARK as alternatives. Bracken extends Kraken-II by combining its taxonomic identification results with Bayesian priors to obtain profiles. For both CLARK and Bracken, we estimated abundance profiles with their default parameters.

On CAMI-II queries, we evaluated CONSULT-II against a host of methods studied by CAMI-II. In particular, we focused on the ten-sample marine dataset (5Gbp each) and the 100-sample (2Gbp each) strain-madness dataset. For alternative methods, submitted results were available from CAMI-II. To make comparisons fair, a new CONSULT-II library was constructed using the reference genomes provided under the scope of the challenge (NCBI RefSeq snapshot as of 2019/01/08); among these, we randomly selected 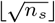 genomes per each species *s* represented by *n*_*s*_ genome (for a total of 18,381 genomes) and included these in the library. We followed the same evaluation approach as the original paper and compared tools by measuring purity versus completeness and L1 norm versus weighted UniFrac error (Lozupone and Knight, 2005). Note that we only included CAMI-II versions of the tools considered, and omitted earlier versions.

## Results

### Empirical evaluation of CONSULT-II heuristics

#### Accuracy of *k*-mer matches

We start by evaluating LCAs of matched *k*-mers across different ranks and HD values (Fig. 2a), keeping in mind that incorrect *k*-mer-level matches do not immediately lead to read-level errors. For the queries with low novelty (MinGND *<* 0.06), ≈9% of query *k*-mers exactly match the reference and have the correct species-level ID; far fewer exact-matches have a soft LCA label at higher taxonomic levels, and the proportion of true matches decreases rapidly with higher HD at all ranks. In contrast, less than 0.5% of the *k*-mers of novel queries (MinGND ≥ 0.22) match any reference, and correct matches peak at HD=1 or HD=2. For mildly novel genomes (0.06 ≤ MinGND *<* 0.22), non-exact matches provide a substantial portion of all correct matches. Unlike true matches, the proportion of false matches tends to increase or remain flat as the HD increases. Nevertheless, in many cases especially at higher taxonomic levels, there are more correct inexact matches with HD ≥ 1 than incorrect ones. For example, 80% of phylum-level inexact matches in the middle novelty range are correct.

**Fig. 2:**
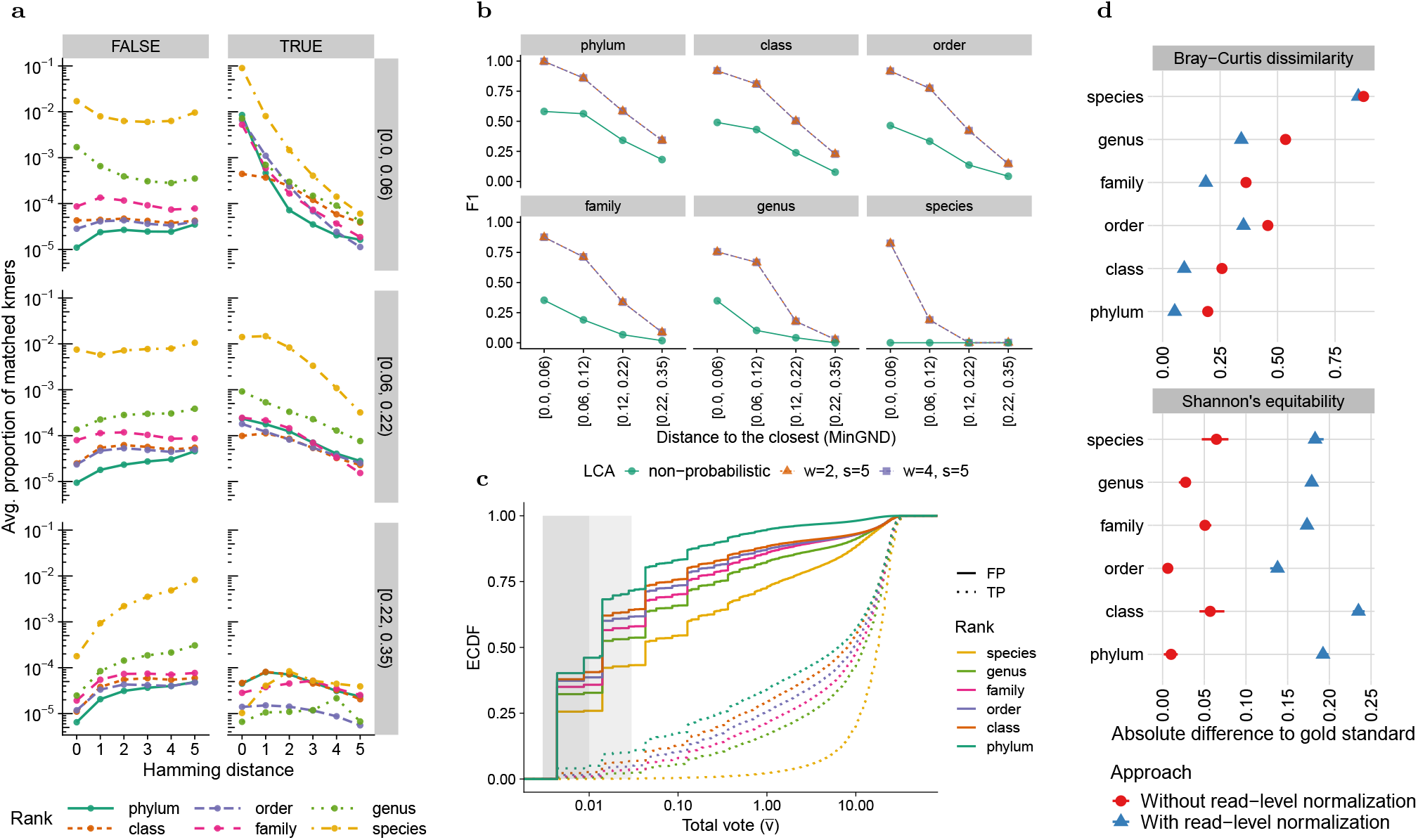
Empirical analysis of LCA update probability *p*_*u*_ and total vote values v (t) across different ranks. (a) The proportion of matched *k*-mers across different ranks, novelty bins (box rows, labeled by MinGND range), and Hamming distances. A match is TRUE if the LCA taxon of the matched *k*-mer is from the correct lineage, and matches are only counted for the rank of the LCA taxon. Note that a FALSE *k*-mer match at a particular rank can be correct at higher ranks but is not counted there. The *y*-axis is the ratio between *k*-mer matches of each type and all 119 × 66667 *k*-mers of a query, averaged over all query genomes. (b) Comparison of soft and hard LCA, based on F1 across different ranks and bins. We compare two parameter settings for the *p*_*u*_ function of soft LCA. (c) The empirical cumulative distribution function (ECDF) of total vote values for correct (TP) and incorrect (FP) classifications, revealing the predictive power of votes. The shaded areas note 0.01 and 0.03 thresholds, used to filter out spurious classifications. (d) Comparison of two different modes of profiling according to Bray-Curtis dissimilarity and Shannon’s equitability. The two measures are computed independently for CONSULT-II’s report for each taxonomic rank. Although the default mode (with read-level normalization, using Eq. (6)) predicts the actual composition more accurately, the optional mode (without read-level normalization) is a better estimator of Shannon’s equitability.

#### Advantages of the soft LCA approach

We next compare the use of hard and soft LCA (Fig. 2b). Overall, soft LCA provides a dramatic improvement compared to naively computing LCA. The improvements are especially large for less novel queries (*<* 0.12 MinGND). Interestingly, at the species rank, hard LCA completely fails while soft LCA has acceptable accuracy for the less novel queries. While a soft LCA approach is clearly helpful, the choice of the ideal probability function is unclear. Since tuning parameters *w* and *s* with a validation set is not practically feasible, we only tested the sensitivity of CONSULT-II performance; settings *w*=2 and *w*=4 are almost identical, with a negligible advantage for *w*=4 (no more than 0.001 in terms of F1). Hence, we use *w*=4 and set it as the default value.

#### Impact of the total vote on classification

Our majority vote mechanism classifies a read at some level, even if the total vote in Eq. (4) is small. However, the total vote correlates with whether a classification is correct (Fig. 2c). In particular, about 35% of the FP predictions have 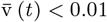 (corresponding to up to two *k*-mers with HD=5), compared to ≈2% of TP predictions. Similarly, around 55% of FPs have 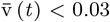 (i.e., less than two matches of HD=4), compared to ≈6% of TPs. Thus, filtering reads with low 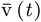 reduces the FPs at the expense of removing some TPs, improving precision and reducing recall (Fig. S3b) with only small changes in F1 scores (Fig. S3a). To ensure our precision levels are similar to other methods, we avoid classifying reads with low total votes (default: 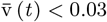).

#### Impact of normalization and unclassified on profiling

Since the total vote values at the root vary widely across reads, we face a question in profiling samples: Should each read contribute equally to the abundance profile of a given sample? If so, we should normalize total vote values per read, as in Eq. (5). Alternatively, we may skip this step and use the total vote values directly in Eq. (6). In CONSULT-II v0.1.1 described in Şapcı *et al*. (2023), we followed the latter approach and skipped Eq. (5); we also used a slightly different equation for Eq. (6) and took the square root of v (*t* ) (Section S1.2). Empirically, adding read-level normalization results in a dramatic improvement in accuracy (Fig. 2d) measured by the Bray-Curtis dissimilarity (up to ≈20%). Surprisingly, skipping the read normalization provides extremely accurate estimates of Shannon’s equitability. Thus, we keep the non-normalized version as a non-default option (since v0.3.0). The use of genome sizes (Eq. (6) versus Eq. (7)) also improves the profile accuracy (Fig. S9). When we added the *unclassified* option to profiles, as much as 35% at the species rank and as little as 6% at the phylum rank were unclassified (Fig. S8a). Adding *unclassified* taxa results in slightly more accurate relative abundances in terms of Bray-Curtis dissimilarity (Fig. S8b). Thus, we include unclassified taxa in output profiles by default (since v0.4.0). Finally, note that Eq. (6) reports independent profiles for each rank. Şapcı et al. (2023) instead reported metrics computed at the species level and aggregated to higher ranks (Fig. S7).

### Comparison to other methods

#### Controlled novelty experiments

On the bacterial query set, CONSULT-II has better F1 scores than CLARK and Kraken-II on all levels above species (Fig. 3a). Only for queries with almost identical genomes in the reference set do all methods have high accuracy with a slight advantage at upper ranks for CONSULT-II. As queries become more novel, accuracy drops across all ranks for all methods. However, CONSULT-II degrades much slower and shows clear improvements for novel genomes. For queries with MinGND *>* 0.05, CONSULT-II outperforms Kraken-II and CLARK across all levels above species with substantial margins. Moreover, the improvements become more substantial at higher taxonomic levels. For instance, with MinGND *>* 0.15, CONSULT-II has mean F1 scores that are 0.12, 0.13, 0.14, and 0.19 more than the second-best method (Kraken-II) respectively for family, order, class, and phylum ranks. Similar patterns are observed when CONSULT-II is run with different total vote thresholds (Fig. S3a). Between CLARK and Kraken-II, Kraken-II has a slight advantage in all levels, except at the species level.

**Fig. 3:**
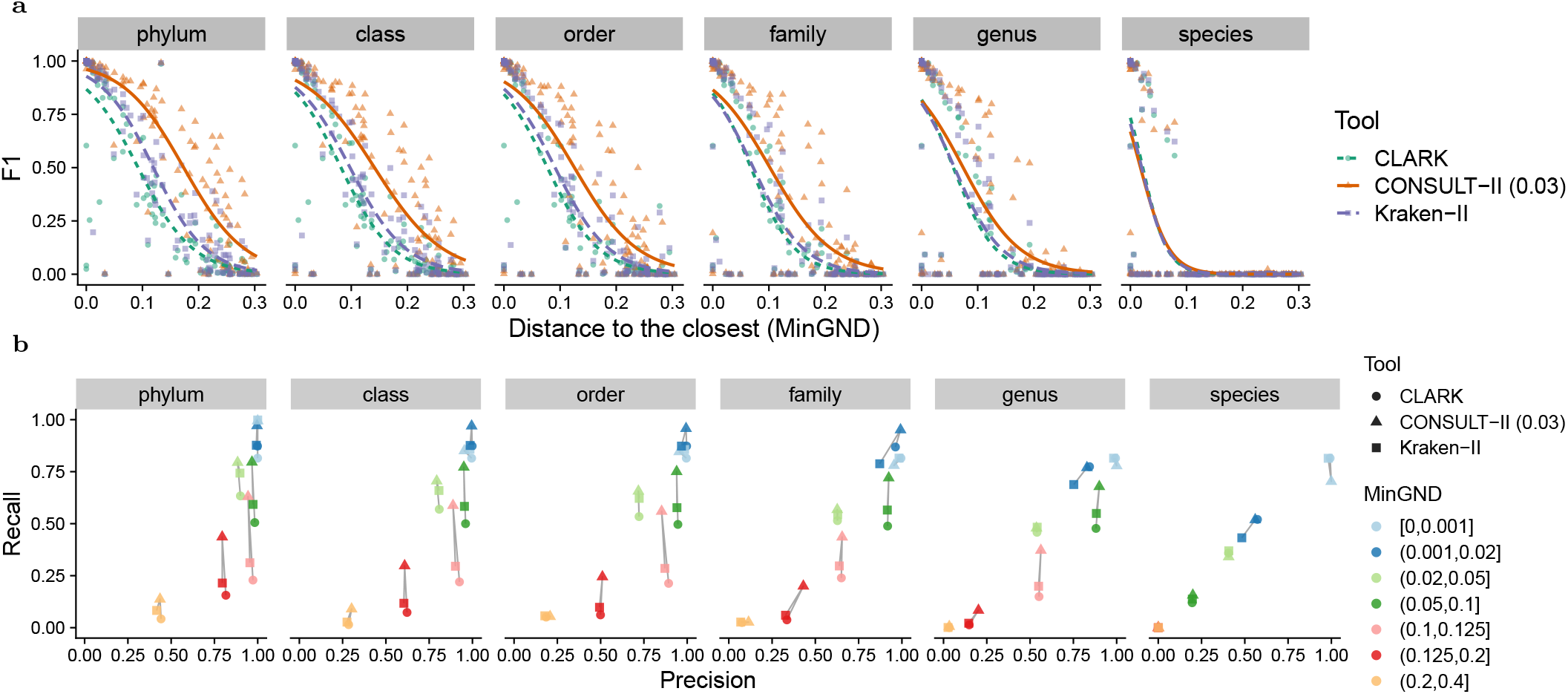
Comparison of CONSULT-II with Kraken-II and CLARK on the controlled novelty experiment with 120 bacterial query genomes and 66667 reads per genome. (a) Accuracy measured using F1 (^2*T P*^*/*(2*T P* + *F P* + *F N* )), showing one dot per query and fitting splines using a generalized linear model. (b) Recall (^*T P*^*/*(*T P* + *F N* )) versus precision (^*T P*^*/*(*T P* + *F P* )). Query genomes are binned into 7 groups based on their Mash-estimated MinGND. Lines connect results for the same bin for ease of comparison.

The advantage of CONSULT-II is due to higher recall and not precision (Fig. 3b) as the precision levels of methods are comparable in most cases, with only slight differences. At the species level, all methods fail to classify moderately novel queries (MinGND *>* 0.12), and CONSULT-II does not show consistent improvements. At the higher levels, the advantage of CONSULT-II is most clear for novel queries which have the closest reference genome within the [0.06, 0.22) MinGND range. For these queries, CONSULT-II often has equal precision to other methods but much higher recall. For the most novel queries (≥ 0.22 MinGND) while CONSULT-II shows some improvement, its precision and recall are still not high. Note that the improved recall of CONSULT-IIfurther increases if we decrease the total vote threshold at the expense of precision (see Fig. S3b).

Results on the archaeal queries are similar to bacterial queries, with some notable differences (Fig. S4). Compared to bacterial queries, all methods tend to have higher F1 scores for queries in the [0.05, 0.2) MinGND range. For all methods, beyond the species rank, the precision tends to be higher regardless of the novelty and recall tends to be lower compared to bacterial queries, especially for less novel genomes. Here, for the bin with the most novel genomes, CONSULT-II has noticeably higher recall and lower precision than alternatives; the gain in the recall offsets the loss in the precision judging by F1.

#### Profiling results for CAMI-I and CAMI-II challenges

For the CAMI-I challenge, CONSULT-II abundance profiles are consistently better than Bracken and CLARK in terms of the Bray-Curtis score (Fig. 4a). At the species level, all methods have high errors and are comparable. CLARK and Kraken-II are similar across ranks, and the advantage of CONSULT-II is most pronounced at higher levels; the second-best method’s error is 40%, 18%, 44%, and 44% higher compared to CONSULT-II at the family to phylum levels, respectively. In terms of Shannon’s equitability, which is a measure of the variety and distribution of taxa present in a sample, all methods underestimate it. CONSULT-II and Bracken are comparable, and they both outperform CLARK substantially. Bracken is closer to the gold standard at the phylum level, while CONSULT-II is better at family and below (Fig. 4a).

**Fig. 4:**
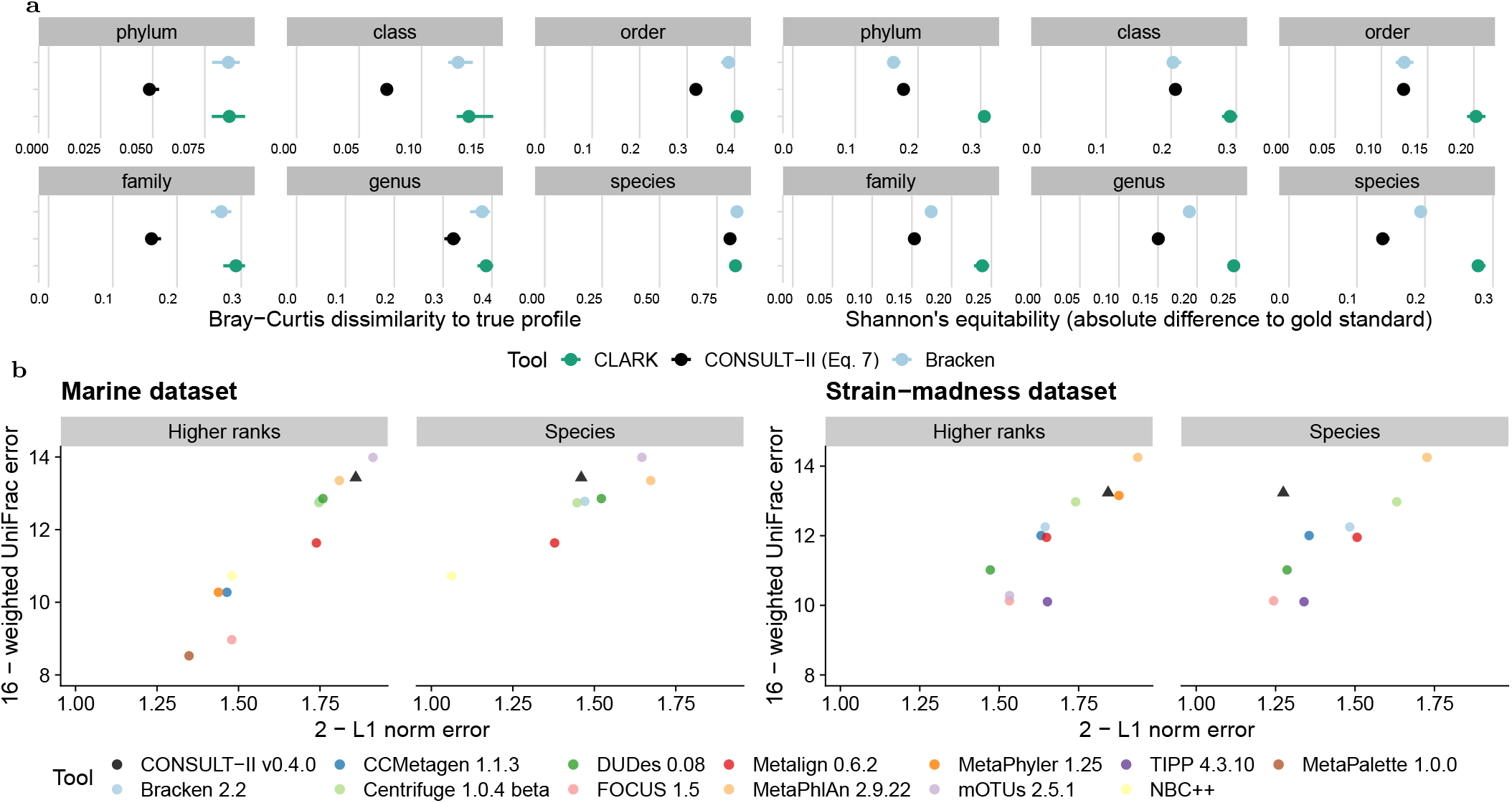
(a) Results on abundance profiling experiments on all five samples of the high-complexity CAMI-I dataset, comparing CONSULT-II with Bracken and CLARK across different taxonomic ranks, using custom libraries constructed with WoL reference database. Left: Bray-Curtis dissimilarity measures the similarity between estimated and true profiles. The smaller the value, the more similar the distributions of abundance profiles are. Right: Shannon’s equitability as a measure of alpha diversity. The closer Shannon’s equitability of the predicted profile to the gold standard, the better it reflects the actual alpha diversity in the gold standard in terms of the evenness of the taxa abundances. The points are mean values across five samples, and the horizontal bars are for standard error. (b) Performance comparison on CAMI-II benchmarking challenge, marine and strain-madness datasets. Following the original paper, we show the upper bound of L1 norm (2) minus actual L1 norm versus the upper bound of weighted UniFrac error (16) minus actual weighted UniFrac error for genus and phylum ranks. Each data point is the average value over 10 marine samples and 100 strain-madness samples. Higher-ranks is mean over all ranks other than species. Metrics are computed using OPAL with default settings and `-n`. For clarity, axes are cut; see Fig. S6 for full results, including all ranks.

On the CAMI-II datasets, we compare across 12 methods (Figs. 4b and S6). CONSULT-II is ranked as the second-best performing method for *both* marine and strain-madness datasets, according to rank-invariant weighted UniFrac error, losing to MetaPhlAn 2.9.22 in the strain-madness dataset and to mOTUs 2.5.1 in the marine dataset (Fig. 4b). Note that MetaPhlAn is ranked 3rd on the marine dataset and mOTUs is ranked 9th in the strain-madness dataset (51% higher weighted UniFrac error than CONSULT-II). CONSULT-II is among the top three tools according to L1 norm error in most cases, except at the species rank (Fig. S6). For species, CONSULT-II did not have high purity and was not among the best for the L1 norm error, especially in the strain-madness dataset. On the marine data, CONSULT-II followed mOTUs, with 0.20 versus 0.13 L1 norm error on average across all ranks. On the strain-madness dataset, it ranked 3rd across all ranks after MetaPhyler and MetaPhlAn with 0.16 versus 0.12 and 0.06 average L1 norm errors across ranks above species (as MetaPhyler lacks species-level profile), respectively.

#### Resource usages of tools

We benchmark resource usage of all tools over queries selected from 30 genomes generated by simulating 66667 short reads from each. CONSULT-II and Kraken-II are more than ×4 and ×9 faster than CLARK, respectively (Fig. 5). While Kraken-II is considerably faster than CONSULT-II (91 vs. 390 seconds), the difference is mostly because CONSULT-II splits the task into two independent stages for the sake of backward compatibility with CONSULT. One subprogram finds *k*-mer matches and writes them to the disk; a second subprogram reads results and performs prediction.

**Fig. 5:**
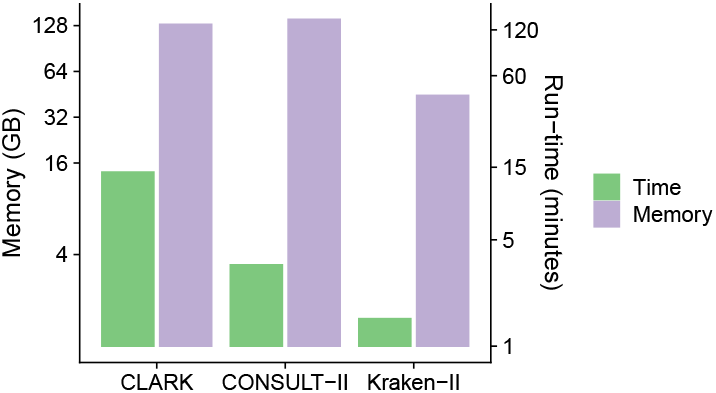
Running time and peak memory usage for 30 queries, each with 66667 150bp reads. Computations used 24 threads, performed on a machine with 2.2 GH_z_ AMD EPYC 7742 processors.

We also observed significant differences in terms of library construction times. Kraken-II was again the fastest tool (≈3.5 hours with 96 threads), followed by CLARK (≈10 hours with 96 threads). CONSULT-II took ≈28 hours (with 128 threads) to construct, and this time was dominated by the soft LCA computation step. Note, however, that library construction is a one-time operation; once constructed, libraries can be distributed.

CONSULT-II, in its default mode, has the highest memory footprint (140Gb), followed by CLARK (130Gb). Kraken-II has much better memory efficiency, using only 44Gb. The CLARK’s memory requirement increased substantially during the library building, exceeding 350Gb. Although it is not as dramatic as CLARK, both CONSULT-II and Kraken-II consumed more memory during the library building (16Gb and 1Gb, respectively).

Since CONSULT-II uses more than three times as much memory as Kraken-II, we asked if its improved performance was because of its higher memory usage. To answer this question, we analyzed the performance of CONSULT-II with a much smaller library (32Gb) by setting *h*=14 and *b*=10 to store fewerl*k*-mers (see Fig. S5). Compared to the default CONSULT library, the lightweight CONSULT-II had 13%, 12%, 5%, 11%, 10%, and 4% decrease in F1, for phylum to species ranks. Nevertheless, compared to Kraken-II, CONSULT-II achieved 16%, 14%, 10%, 11%, 5% higher F1 scores for phylum to genus ranks, and 6% less at species, despite using less memory (32Gb versus 44Gb). Thus, the advantages of CONSULT-II over Kraken-II persist, especially for higher ranks, but at a lower level when using similar memory.

## Discussion

We introduced CONSULT-II for taxonomic classification and abundance profiling. Our approach uses LSH to efficiently find inexact *k*-mer matches and the match distances between reference genomes and a query. Heuristics are then used to translate *k*-mer-level matches and their HD to read-level classification and sample-level profiling. While they lack theoretical guarantees, these heuristics performed remarkably well in our experiments and outperformed popular *k*-mer-based methods. In particular, our equations for LCA update probability (2) and vote-versus-distance (3) are based on intuitive assumptions and expectations, without much fine-tuning and very few parameters. Future research should explore alternative approaches, including using machine learning to automatically train parameter-rich functions instead of heuristics. To ensure robustness, it would be essential to evaluate such fine-tuned methods on more varied datasets.

An alternative direction of future work is developing a theoretical framework for translating *k*-mer distances to taxonomic classifications. Connecting taxonomic profiling to distance-based phylogenetic placement could provide a framework to tackle this goal. Such a framework may allow us to go beyond taxonomic identification and could provide alignment-free phylogenetic placement of reads, as others have attempted (Blanke and Morgenstern, 2020). Finally, future work should address the high memory consumption of CONSULT-II compared to alternatives, perhaps by smart subsampling of *k*-mers.

## References

Ames, S. K. et al. (2013). Scalable metagenomic taxonomy classification using a reference genome database. Bioinformatics (Oxford, England), 29(18), 2253–2260.

Asnicar, F. et al. (2020). Precise phylogenetic analysis of microbial isolates and genomes from metagenomes using PhyloPhlAn 3.0. Nature Communications, 11(1), 2500. Publisher: Springer US ISBN: 0348556977.

Berlin, K. et al. (2015). Assembling large genomes with singlemolecule sequencing and locality-sensitive hashing. Nature biotechnology, 33(6), 623–630.

Blanke, M. and Morgenstern, B. (2020). Phylogenetic placement of short reads without sequence alignment. bioRxiv, pages 2020–10. Publisher: Cold Spring Harbor Laboratory.

Brown, D. and Truszkowski, J. (2013). LSHPlace: Fast phylogenetic placement using locality-sensitive hashing. In Pacific Symposium On Biocomputing, pages 310–319. ISSN: 2335-6936.

Buhler, J. (2001). Efficient large-scale sequence comparison by locality-sensitive hashing. Bioinformatics, 17(5), 419–428.

Choi, J. et al. (2017). Strategies to improve reference databases for soil microbiomes. The ISME Journal, 11(4), 829–834.

Handelsman, J. (2004). Metagenomics: application of genomics to uncultured microorganisms. Microbiology and molecular biology reviews, 68(4), 669–85.

Har-Peled, S. et al. (2012). Approximate nearest neighbors: Towards removing the curse of dimensionality. Theory of Computing, 8(1), 321–350.

Huang, W. et al. (2012). ART: A next-generation sequencing read simulator. Bioinformatics, 28(4), 593–594.

Lau, A.-K. et al. (2019). Read-SpaM: assembly-free and alignment-free comparison of bacterial genomes with low sequencing coverage. BMC Bioinformatics, 20(S20), 638.

Liang, Q. et al. (2020). DeepMicrobes: taxonomic classification for metagenomics with deep learning. NAR Genomics and Bioinformatics, 2(1).

Liu, B. et al. (2011). MetaPhyler: Taxonomic profiling for metagenomic sequences. In Bioinformatics and Biomedicine (BIBM), 2010 IEEE International Conference on, pages 95–100. IEEE.

Locey, K. J. and Lennon, J. T. (2016). Scaling laws predict global microbial diversity. Proceedings of the National Academy of Sciences, 113(21), 5970–5975.

Lozupone, C. and Knight, R. (2005). UniFrac: a new phylogenetic method for comparing microbial communities. Applied and environmental microbiology, 71(12), 8228–8235.

Lu, J. et al. (2017). Bracken: estimating species abundance in metagenomics data. PeerJ Computer Science, 3, e104.

Luo, Y. et al. (2019). Metagenomic binning through low-density hashing. Bioinformatics, 35(2), 219–226.

McDonald, D. et al. (2023). Greengenes2 unifies microbial data in a single reference tree. Nature Biotechnology, pages 1–4. Publisher: Nature Publishing Group.

McIntyre, A. B. R. et al. (2017). Comprehensive benchmarking and ensemble approaches for metagenomic classifiers. Genome Biology, 18(1), 182.

Meyer, F. et al. (2019). Assessing taxonomic metagenome profilers with OPAL. Genome Biology, 20(1), 51.

Meyer, F. et al. (2022). Critical Assessment of Metagenome Interpretation: the second round of challenges. Nature Methods, 19(4), 429–440.

Milanese, A. et al. (2019). Microbial abundance, activity and population genomic profiling with mOTUs2. Nature Communications, 10(1), 1014.

Nasko, D. J. et al. (2018). RefSeq database growth influences the accuracy of k-mer-based lowest common ancestor species identification. Genome Biology, 19(1), 165.

Ondov, B. D. et al. (2016). Mash: fast genome and metagenome distance estimation using MinHash. Genome Biology, 17(1), 132.

Ounit, R. et al. (2015). CLARK: fast and accurate classification of metagenomic and genomic sequences using discriminative kmers. BMC Genomics, 16(1), 236.

Pachiadaki, M. G. et al. (2019). Charting the Complexity of the Marine Microbiome through Single-Cell Genomics. Cell, 179(7), 1623–1635.e11.

Parks, D. H. et al. (2020). A complete domain-to-species taxonomy for Bacteria and Archaea. Nature Biotechnology, 38(9), 1079–1086.

Rachtman, E. et al. (2020). The impact of contaminants on the accuracy of genome skimming and the effectiveness of exclusion read filters. Molecular Ecology Resources, 20(3), 1755–0998.13135.

Rachtman, E. et al. (2021). CONSULT: accurate contamination removal using locality-sensitive hashing. NAR Genomics and Bioinformatics, 3(3), 10.1101/2021.03.18.436035.

Rasheed, Z. et al. (2013). 16S rRNA metagenome clustering and diversity estimation using locality sensitive hashing. BMC Systems Biology, 7(Suppl 4), S11.

Şapcı, A. O. B. et al./person-group>. (2023). Consult-ii: Taxonomic identification using locality sensitive hashing. In K. Jahn and T. Vinař, editors, Comparative Genomics, pages 196–214, Cham. Springer Nature Switzerland.

Sczyrba, A. et al. (2017). Critical Assessment of Metagenome Interpretation—a benchmark of metagenomics software. Nature Methods, 14(11), 1063–1071.

Segata, N. et al. (2012). Metagenomic microbial community profiling using unique clade-specific marker genes. Nature Methods, 9(8), 811–814. ISBN: 1548-7091.

Shah, N. et al. (2021). TIPP2: metagenomic taxonomic profiling using phylogenetic markers. Bioinformatics, 37(13), 1839–1845.

Sunagawa, S. et al. (2013). Metagenomic species profiling using universal phylogenetic marker genes. Nature Methods, 10(12), 1196–1199.

Truong, D. T. et al. (2015). MetaPhlAn2 for enhanced metagenomic taxonomic profiling. Nature Methods, 12(10), 902–903.

von Meijenfeldt, F. A. B. et al. (2019). Robust taxonomic classification of uncharted microbial sequences and bins with CAT and BAT. Genome Biology, 20(1), 217.

Wood, D. E. et al. (2019). Improved metagenomic analysis with Kraken 2. Genome Biology, 20(1), 257.

Wu, S. et al. (2020). GMrepo: a database of curated and consistently annotated human gut metagenomes. Nucleic Acids Research, 48(D1), D545–D553.

Ye, S. H. et al. (2019). Benchmarking Metagenomics Tools for Taxonomic Classification. Cell, 178(4), 779–794.

Zhu, Q. et al. (2019). Phylogenomics of 10,575 genomes reveals evolutionary proximity between domains Bacteria and Archaea. Nature Communications, 10(1), 5477.

Zhu, Q. et al. (2022). Phylogeny-Aware Analysis of Metagenome Community Ecology Based on Matched Reference Genomes while Bypassing Taxonomy. mSystems, 7(2), e0016722.

