## Supplementary Materials for "CONSULT-II: Accurate taxonomic identification and profiling using locality-sensitive hashing"

### Supplementary Materials for "CONSULT-II: Taxonomic identification using locality-sensitive hashing"

### S1 Supplementary Notes

#### S1.1 Automatic adjustment of the hash table size

We compute appropriate $h$ and $l$ values to accommodate $G$ $k$-mers, and to maintain the same performance for a fixed $b$. Each table can store $2^{2h}b$ $k$-mers, and thus, we need $h = {}^1\!/_2 \log_2 ({}^G\!/_b)$ to accommodate $G$ $k$-mers in all $l$ tables. Being able to control the value of $\rho(d) = \alpha$ allows us to ensure the desired level of performance. Solving for $l$ in $\rho(d) = 1 - (1 - (1 - \frac{d}{k})^h)^l$ yields $l = {}^{\log(1-\alpha)}\!/_{\log\left(1-(1-p/k)^h\right)}$. Thus, for a given $b$, our heuristic can select $h$ and $l$ to accommodate all $k$-mers, and fix the performance (Fig. S10a). Remarkably, $l$ only ranges between 1 and 3 for even very large or small $G$. Then, to choose $b$, we simply search a reasonable discrete range for the value that minimizes the total memory consumption of CONSULT-II, which is given by $4l2^{2h}b + 8G$. Adding some bounds and appropriate rounding, we obtain:

$$h(b) = \max\left\{\left\lceil {}^1\!/_2 \log_2 ({}^G\!/_b)\right\rceil, 4\right\} \tag{1}$$

$$l(b) = \max\left\{\mathrm{round}\left({}^{\log(1-\alpha)}\!/_{\log\left(1-(1-p/k)^h\right)}\right), 2\right\} \tag{2}$$

$$\hat{b} = \underset{4\le b\le 20}{\arg\min}\left(4\cdot l(b)2^{2h(b)}b + 8\cdot G\right). \tag{3}$$

Thus, we set $b = \hat{b}, l = l(\hat{b})$, and $h = h(\hat{b})$. The user can select $\alpha$ and $p$, CONSULT uses $\rho(3)$=0.4 by default. CONSULT-II, slightly increase the level of performance to $\rho(3)$=0.45 (i.e., $\alpha$=0.45 for $p$=3). As indicated above, the right choice of $b$ as described above can lead to substantial memory saving, all else equal (Fig. S10b). Note that the best choice of $b$ depends on $G$ in very non-linear ways due to the necessary rounding in the settings. We compare CONSULT and CONSULT-II on a dataset consisting of six E. coli genomes in the reference set (12 million 32-mers), and four E. coli genomes in the query set (1 million 150bp reads). Fig. S10c demonstrates the dramatic improvement in terms of computational resources obtained. Our heuristic determines the optimal parameter configuration as $h$=10, $b$=12, and $l$=2 and achieves the same level of accuracy with less than 500Mb. As expected, having a smaller library also leads to shorter runtimes, both for library construction and querying. Library construction takes $\approx 60\times$ shorter thanks to CONSULT-II's heuristic. At the query time, CONSULT-II is $\approx 7\times$ faster given a single thread. CONSULT-II is also efficiently parallelized and obtains close to linear speed-ups up to 8 threads.

#### S1.2 Alternative mode of abundance profiling

For the optional profiling mode of CONSULT-II, we skip read-level normalization and aggregate all total votes, $\bar{\mathrm{v}}_{\mathcal{R}_\mathrm{i}}(t)$ for taxon $t \in \mathcal{T}_r$, across all reads, $\mathcal{R}_1, \ldots, \mathcal{R}_n$, together into a profile. Similar to the default mode, we consider each taxonomic rank separately. At a given rank $r$, we normalize the sum of square roots of total votes to obtain a profile whose values sum up to 1. While there are many alternatives to consider, we opted to use the following for its superior performance observed empirically;

$$p_t^r = \frac{\sum_i^n \sqrt{\bar{\mathrm{v}}_{\mathcal{R}_\mathrm{i}}(\mathrm{t})}}{\sum_{t'\in\mathcal{T}_r}\sum_i^n \sqrt{\bar{\mathrm{v}}_{\mathcal{R}_\mathrm{i}}(\mathrm{t'})}}$$

where $p_t^r$ is the profile value of taxon $t$ from rank $r$. Then, the abundance profile for rank $r$ is given by $\mathsf{p}^r = \left[p_t^r\right]_{t\in\mathcal{T}_r}$.

### S2 Exact commands and versions

Here, we provide the exact commands that we used to run external tools & CONSULT-II's software throughout our experiments. We also provide version information for reproducibility.

#### S2.1 Computation of genomic distances with Mash [7]

We used Mash (version `1.1`) to estimate genomic distances, using the following command.

```
mash dist FASTQ_FILE_ONE FASTQ_FILE_TWO
```

#### S2.2 Short read simulation with reference using ART [1]

We simulated short reads with length $L = 150$ and coverage $c$, with default error and quality profiles of Illumina HiSeq 2500 using ART (version `2.5.8` – single read mode) with the command below.

```
art_illumina -ss HS25 -i INPUT_FASTA_FILE -l 150 -f c -na -s 10 -o OUTPUT_FASTQ_FILE
```

#### S2.3 Extraction of $k$-mer sets using Jellyfish [4]

We used Jellyfish (version `2.3.0`) to compute unique $k$-mer sets and $k$-mer profiles (for 35bp and 32bp canonical $k$-mers) of reference genomes (FASTA). The command is given below.

```
jellyfish count -m KMER_LENGHT -s 100M -t 24 -C INPUT_FASTA_FILE -o COUNT_JF_FILE
```

Then, to output the list of distinct $k$-mers together with their counts into a FASTQ file, we used the following command. Note that we do not use count values.

```
jellyfish dump COUNT_JF_FILE > OUTPUT_FASTA_FILE
```

#### S2.4 Downsampling reads with `seqtk` [2]

To subsample read collection down to a specified number of reads, denoted by $n$ here, we used `seqtk` (version `1.3r106`) with the command below.

```
seqtk sample -s150 INPUT_FASTQ_FILE n > OUTPUT_FASTQ_FILE
```

#### S2.5 Minimization of $k$-mer sets

For minimization of canonical $k$-mer sets, we used a custom script (version `v0.3.0`), provided with CONSULT-II's software package. The input to this script is a Jellyfish FASTA file containing 35bp canonical $k$-mers extracted from reference genomes, and it outputs their 32bp minimizers in FASTA format. You can compile with `g++ minimize.cpp -std=c++11 -O3 -o minimize`, and then running the following command would generate 32bp minimizers in FASTA format.

```
./minimize -i CANONICAL_35MERS_FASTA -o MINIMIZER_32MERS_FASTA
```

#### S2.6 Taxonomic identification with CONSULT-II

The version of CONSULT-II we presented in this paper is `v0.4.0`.
We used the command below to construct the reference library:

```
./consult_map -p 3 -t 2 -i COMBINED_FASTA_FILE -o LIBRARY_NAME
```

where `-t` is tag size in bits (it determines the number of partitions) and `-p` is the Hamming distance threshold for a match. After constructing the reference library, we added taxonomic information to the reference library (soft LCA information per $k$-mer) with the following two commands.

```
./consult_search -q REFERENCE_GENOMES_DIR -i LIBRARY_NAME -o OUTPUT_DIR \
--taxonomy-lookup-path TAXONOMY_LOOKUP_FILE --filename-map-path FILENAME_MAP_FILE --init-ID
./consult_search -q REFERENCE_GENOMES_DIR -i LIBRARY_NAME -o OUTPUT_DIR \
--taxonomy-lookup-path TAXONOMY_LOOKUP_FILE --filename-map-path FILENAME_MAP_FILE --update-ID
```

where `REFERENCE_GENOMES_DIR` is a directory in which all reference genomes are stored in FASTQ format. Note that `TAXONOMY_LOOKUP_FILE` is an auxiliary file that can be generated from a NCBI taxonomy file (`nodes.dmp`), using a script we provide, as below.

```
python scripts/construct_taxonomy_lookup.py -from-taxonomy-database /path/to/nodes.dmp
```

Then, in order to query a set of sequences against a reference library, we used the following CONSULT-II command.

```
./consult_search -i LIBRARY_NAME -q QUERY_FASTA_FILE -o OUTPUT_DIR \
--maximum-distance 5 --save-matches --thread-count 24
```

This command outputs match information files per query, with filenames that are query filenames prefixed with `match-info_*`, to `OUTPUT_DIR`.

Finally, we performed taxonomic classification and abundance profile estimation, using CONSULT-II's sub-commands `consult_classify` and `consult_profile` respectively.

```
./consult_classify -i MATCH_INFO_FILE -o OUTPUT_DIR --taxonomy-lookup-path TAXONOMY_LOOKUP_FILE
./consult_profile -i MATCH_INFO_FILE -o OUTPUT_DIR --taxonomy-lookup-path TAXONOMY_LOOKUP_FILE
```

The above command creates a report which contains total vote values before normalization. An additional script is provided for genome size correction and normalization across taxa at each rank.

```
python scripts/postprocess_profiles.py THRESHOLD PROFILE_FILE NCBI_TAXONOMY_DIR METADATA_FILE S_ID
```

Here, THRESHOLD (1000 in our case) is the total vote threshold for a taxon to be included, `PROFILE_FILE` is the CONSULT-II's output, `NCBI_TAXONOMY_DIR` is a directory which stores `nodes.dmp` and `names.dmp` files, `METADATA_FILE` is tab seperated file which is required to include `species_id` and `total_length` columns for each reference genome. The `total_length` corresponds to the genome size of the reference genome belonging to species `species_id`. `S_ID` is a simple sample identifier name to be included in the header of the report.

#### S2.7 Taxonomic identification with Kraken-II [10] and Bracken [3]

We used version `2.0` and version `2.8` for Kraken-II and Bracken, respectively.

##### S2.7.1 Kraken-II

In order to construct a standard Kraken-II reference library, we used the default command.

```
kraken2-build --standard --no-masking --use-ftp --db DATABASE_NAME
```

To build a custom Kraken-II reference library, we used a set of commands, given below.

1. Download the taxonomy.
   `kraken2-build --download-taxonomy --no-masking --use-ftp --db DATABASE_NAME` using `ftp`.
2. Rename all file extensions to `.fa` from `.fna`.
   ```
   find . -name "*.fna" -exec sh -c 'mv "$1" "${1%.fna}.fa"' _ {} \;
   ```
3. Add custom genomes to the reference library.
   ```
   find genomes/ -name "*.fa" -print0 | xargs -0 -I -n1 kraken2-build --no-masking \
   --add-to-library {} --db DATABASE_NAME
   ```
4. Build the database with specified $k$-mer length `KMER_LEN`, minimizer length `MINIMIZER_LEN` and number of wind-carding positions $s$.
   ```
   kraken2-build --build --no-masking --kmer-len KMER_LEN --minimizer-len MINIMIZER_LEN \
   --minimizer-spaces s --use-ftp --db DATABASE_NAME
   ```

To query against the Kraken-II reference library at variable confidence level $\alpha$, we used the command below.

```
kraken2 --use-names --threads 24 --report REPORT_FILE_NAME --db DATABASE_NAME --confidence α \
--classified-out CLASSIFIED_FASTQ_FILE --unclassified-out UNCLASSIFIED_FASTQ_FILE \
QUERY_FASTQ_FILE > KRAKEN_OUTPUT_FILE
```

##### S2.7.2 Bracken

To process a Kraken-II database for Bracken by dividing each genome into read-length $k$-mers and classify each of those $k$-mers, run the following command with 24 threads (`-t`), where `DATABASE_NAME` is a built Kraken-II database.

```
./bracken-build -d DATABASE_NAME -t 24 -k KMER_LEN -l READ_LEN
```

To perform abundance estimation with Bracken, we used the command below, with default parameters.

```
bracken -d DATABASE_NAME -i KRAKEN_OUTPUT_FILE -o BRACKEN_OUTPUT_FILE -r READ_LEN -l S -t 10
```

#### S2.8 Taxonomic identification with CLARK [8]

We used CLARK (version `1.2.6.1`) with its default parameters. CLARK failed to set target IDs for 494 genomes out of 10470 ToL genomes during the construction of its database. These genomes were added manually by setting the taxon. We followed the below steps to build a custom CLARK database of discriminative $k$-mers in the directory `DIR_DB/`.

1. Create the directory `Custom/` inside `DIR_DB/` directory.
2. Copy reference ToL FASTA files with accession numbers into the `Custom/` directory.
3. Run `./set_target.ssh DIR_DB/ custom --species` command.

We queried sequences against the CLARK database using the default mode of classification (i.e., `-m 1`) with the command below.

```
./classify_metagenome.sh -O SAMPLE_FASTQ_FILE -R RESULT_CSV -n 24
```

### S3 Supplementary Figures

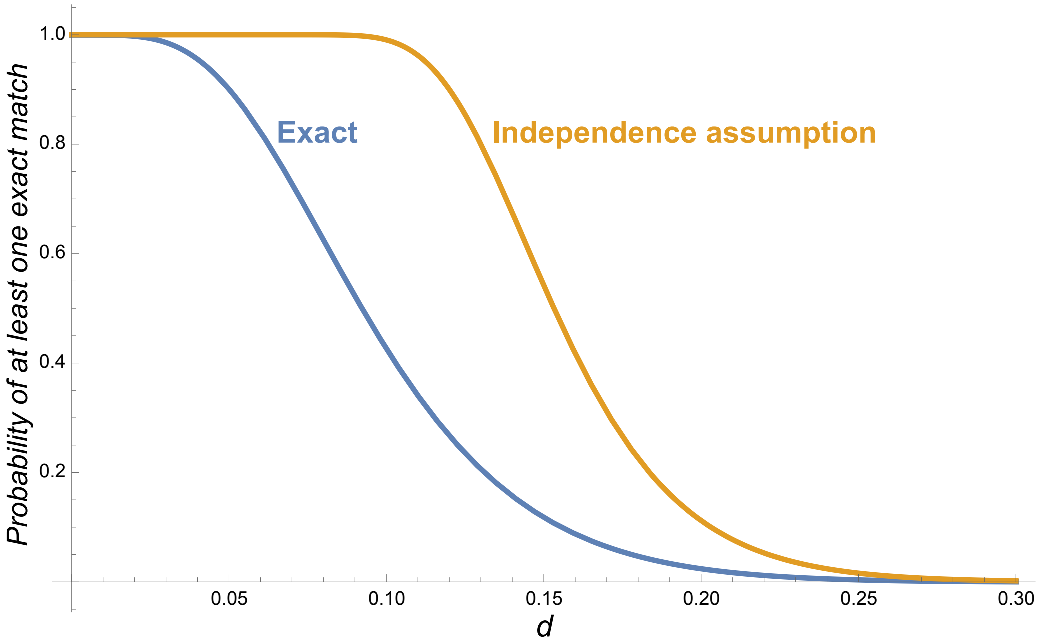


$$p(d, L, k) = \begin{cases} 0 & k > L \\ (1-d)^k & k = l \\ p(d, L-1, k) + (1 - p(k, L-k-1, d))\, d(1-d)^k & k < L \end{cases}$$

$$p_{ind} = 1 - \left(1 - (1-d)^k\right)^{L-k+1}$$

Figure S1: Independence assumption does not work. Probability of at least one exact match $k$-mer ($k$=32) match in a read of length $L$ (=150) at a distance $d$. The exact solution can be derived using dynamic programming recursion shown in blue. The main recursion follows from the observation that the leftmost match is either included in the first $L-1$ positions, or not; if not, the final $k$-mer must match, and the position right to the left of the final kmer should have a mutation. The recursion is computed with memorization using Mathematica. Note the substantial difference from this exact calculation and what would be obtained if we assume independence of $k$-mers.

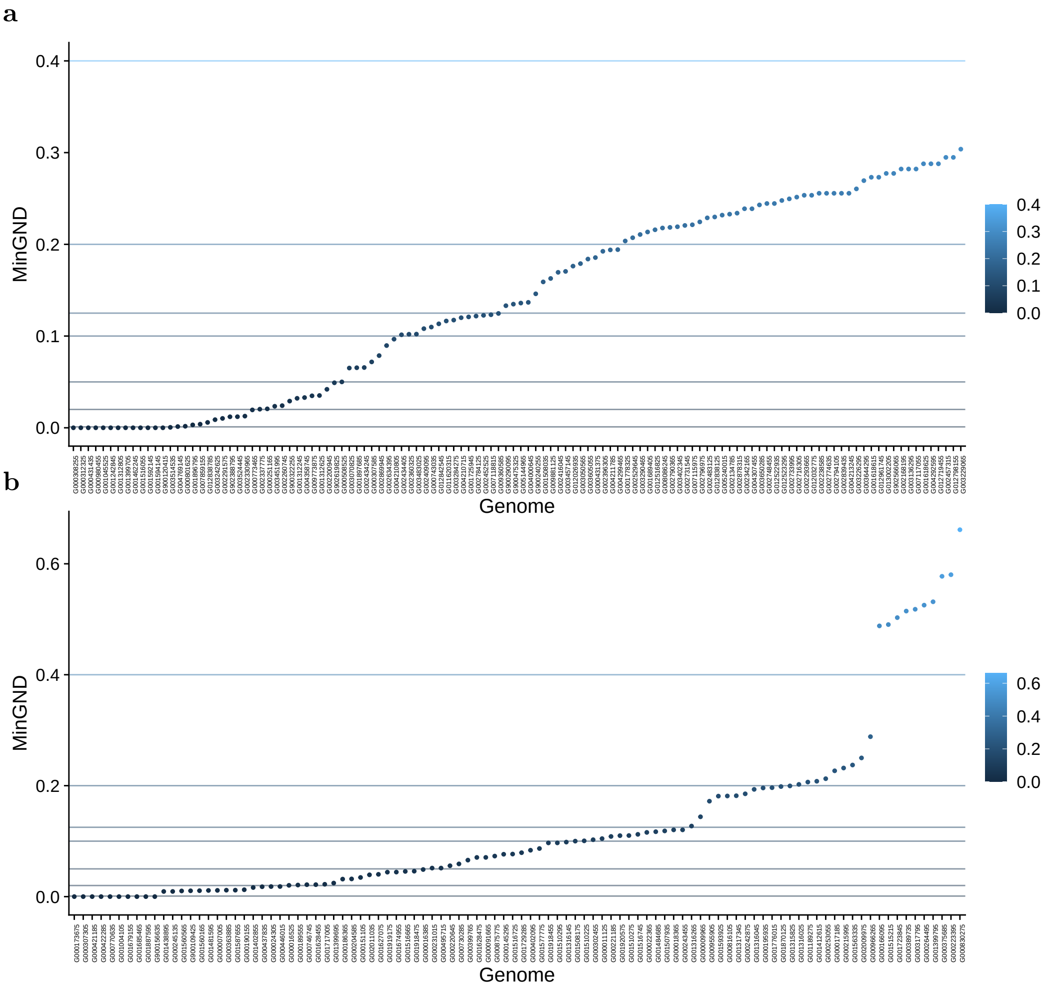


Figure S2: The distribution of distances to the closest genome in the reference set for the (a) 120 bacteria and (b) 100 archaea queries selected. For each query, we show their distance to the closest species, as estimated by Mash [7]. We split queries into seven bins based on these distances (cutting at $0.001, 0.02, 0.06, 0.12, 0.16, 0.22$ and $0.001, 0.02, 0.05, 0.1, 0.125, 0.2, 0.4$) making sure each bin has at least 11 and 10 queries, respectively for bacteria and archaea. Dashed lines show the boundaries between bins of queries by distance.

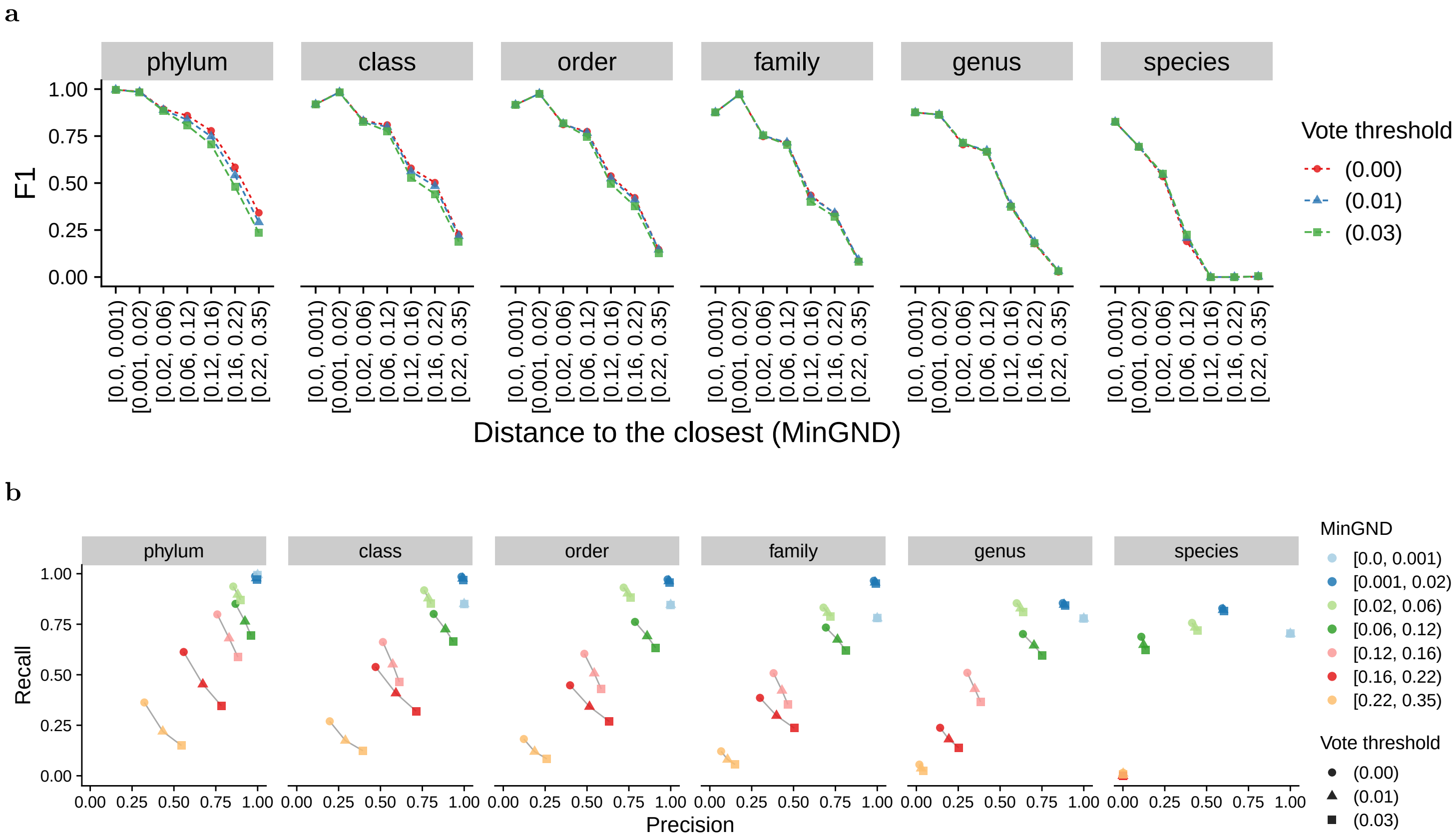


Figure S3: Effect of filtering of predictions performed based on minimum required total vote value on CONSULT-II's performance. (a) F1 score and (b) precision-recall comparison across different bins of distance to the closest reference genome and taxonomic ranks.

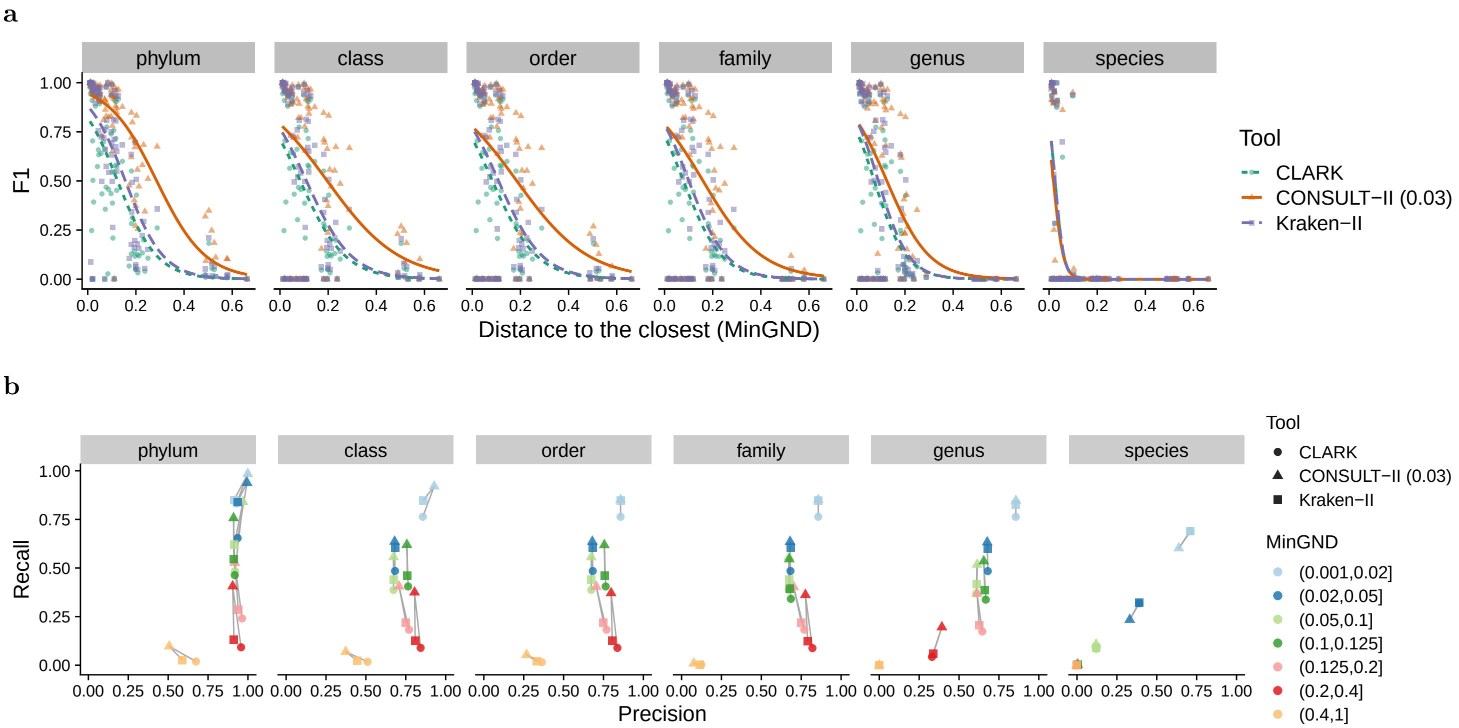


Figure S4: Evaluation of CONSULT-II's classification algorithm in comparison with Kraken-II and CLARK on the controlled novelty experiment with archaea queries. The set of 100 genomes is binned into 7 groups based on their Mash distance to their closest neighbor in the reference set. Results summarize classifications across 66667 reads per query. (a) A fine-grained comparison of the performances of all three methods across different taxonomic levels using F1 ($^{2TP}/_{(2TP+FP+FN)}$) score across taxonomic groups. Each dot stands for a query, and the lines are fitted using a Generalized Linear Model. (b) At different taxonomic levels, recall ($^{TP}/_{(TP+FN)}$) versus precision ($^{TP}/_{(TP+FP)}$) for each bin of queries. The results of different tools from the same bin are connected with a line.

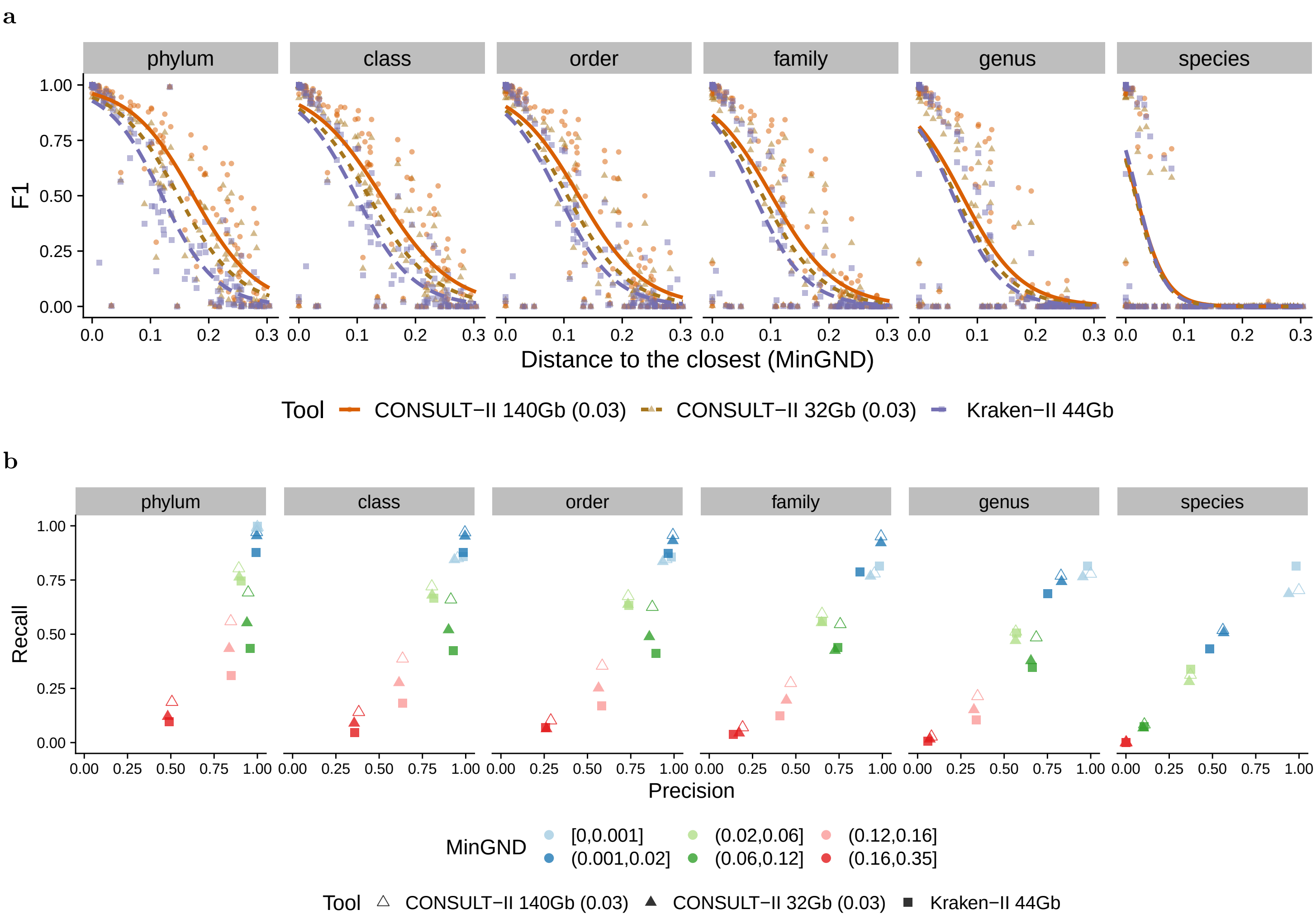


Figure S5: Evaluation of CONSULT-II's classification performance with respect to different library sizes (140Gb and 32Gb), resulting from different $h$ and $b$ configurations, on the controlled novelty experiment with bacteria queries. We set $h{=}15, b{=}7$ and $h{=}14, b{=}10$ respectively for 140Gb and 32Gb libraries. CONSULT-II can achieve higher F1 scores by improving recall over Kraken-II, using less memory (32Gb versus 44Gb). (a) A fine-grained comparison of the performances of all three methods across different taxonomic levels using F1 ($^{2TP}/_{(2TP+FP+FN)}$) score across taxonomic groups. Each dot stands for a query, and the lines are fitted using a Generalized Linear Model. (b) At different taxonomic levels, recall ($^{TP}/_{(TP+FN)}$) versus precision ($^{TP}/_{(TP+FP)}$) for each bin of queries.

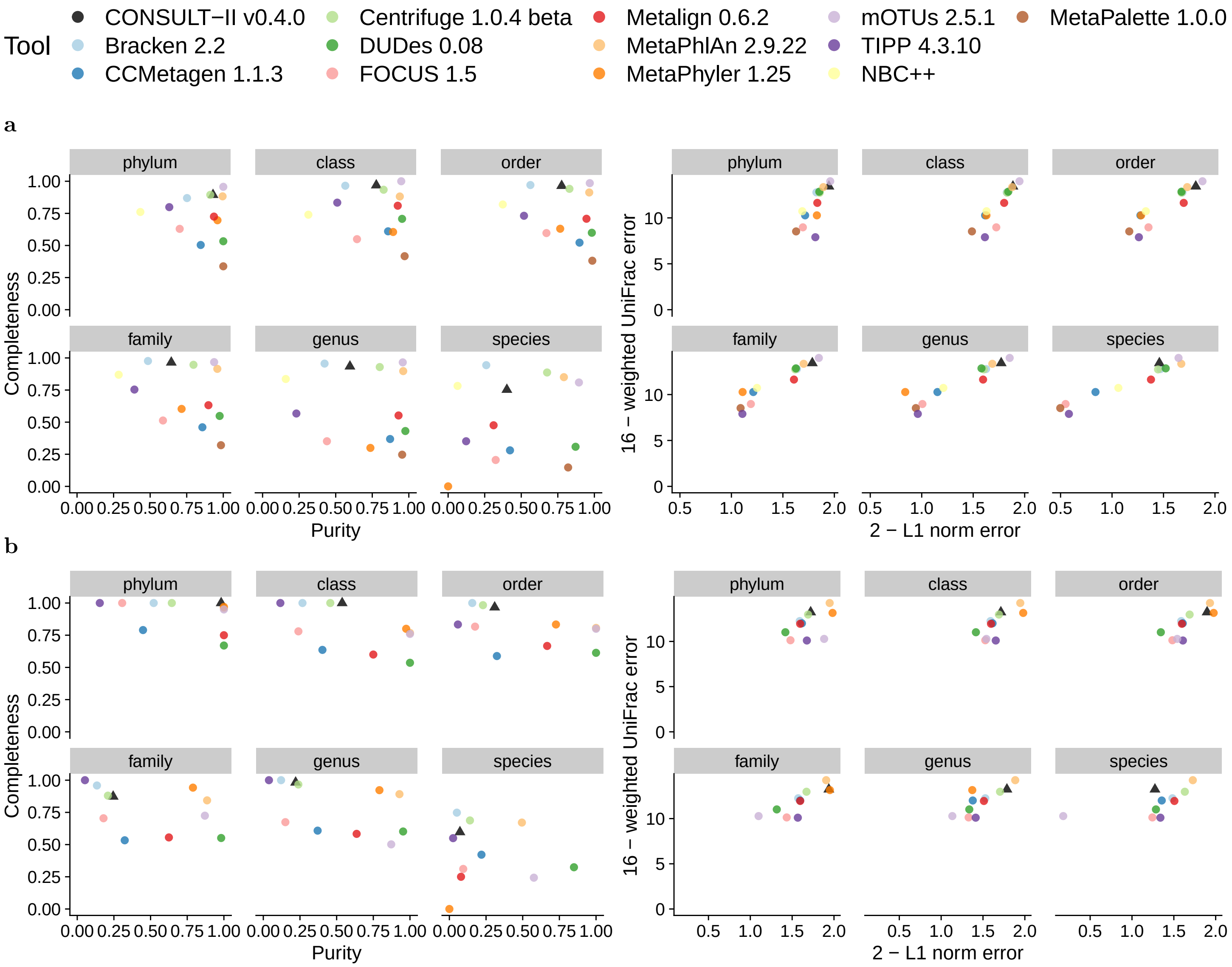


Figure S6: Evaluation of CONSULT-II with CAMI-II[6] benchmarking challenge datasets: (a) marine samples and (b) strain-madness samples. As in the original paper, we show purity versus completeness and the upper bound of L1 norm (2) minus actual L1 norm versus the upper bound of weighted UniFrac error (16) minus actual weighted UniFrac error. Each data point stands for the mean over 10 marine and 100 strain-madness samples. Metrics were determined using OPAL with default settings and the `-n` option.

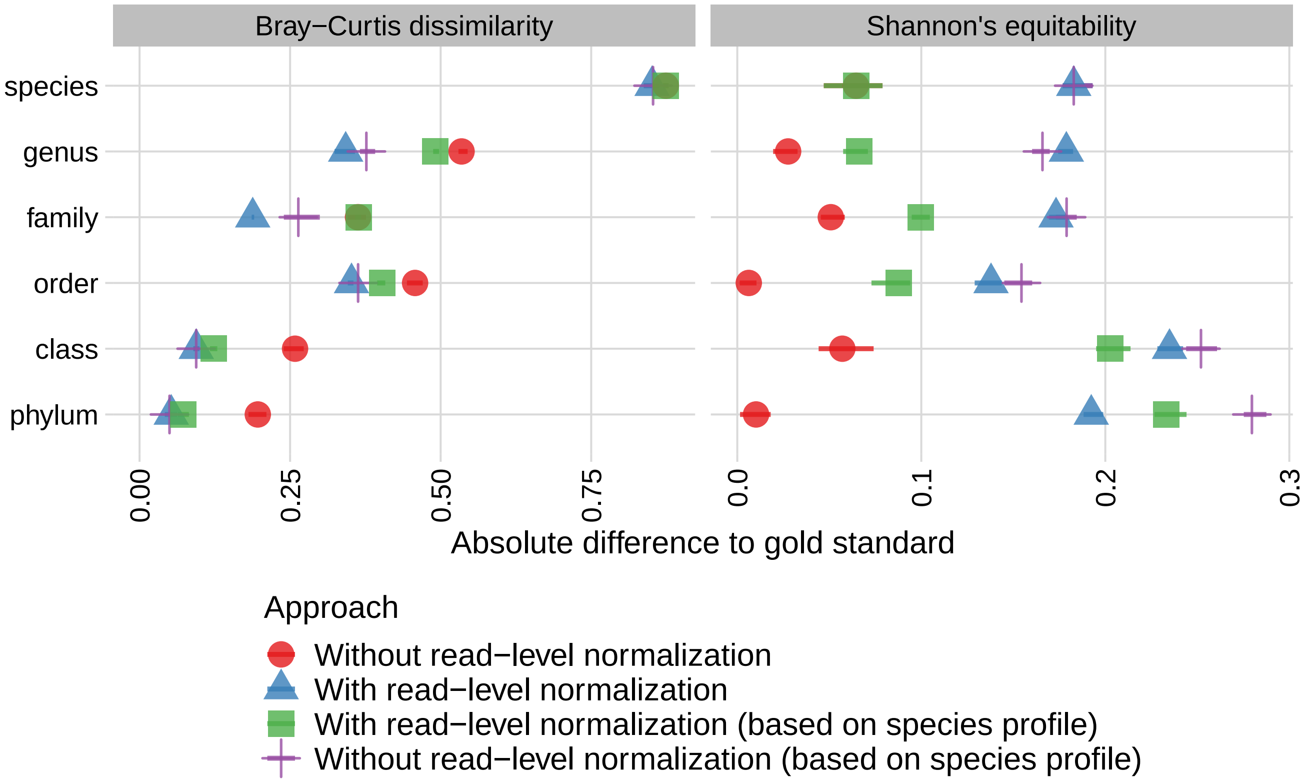


Figure S7: Comparison of two different modes of profiling according to Bray-Curtis dissimilarity and Shannon's equitability. We evaluate the two modes according to measures computed i) based solely on the species-level profile report of CONSULT-II and ii) based on measures computed independently for each rank's profile report. For i), upper taxonomic ranks are derived directly from species-level profile vectors. Bray-Curtis dissimilarity and Shannon's equitability computed according to ii) is what Şapcı et al. [9] reported. In this paper, we switch to a better approach and measure accuracy for each level separately.

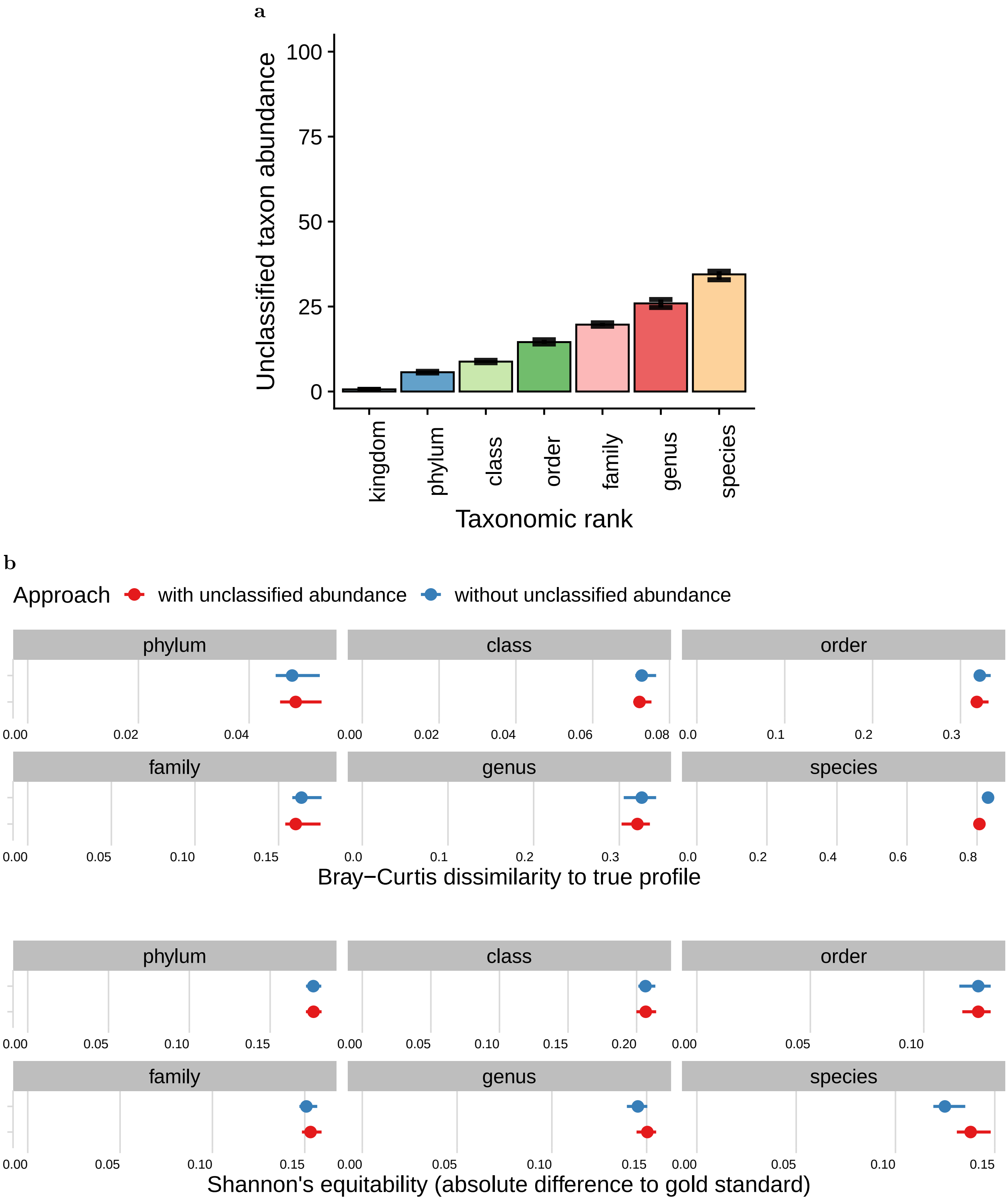


Figure S8: (a) Proportion of relative abundances determined to be *unclassified* across different ranks. (b) Effect of adding artificial nodes for unclassified abundance on the performance. CONSULT-II with unclassified abundance results in more accurate profile estimates across all taxonomic ranks, generally having lower Bray-Curtis dissimilarity to the true profile. Metrics were computed using OPAL[5] with the `-n` option to normalize.

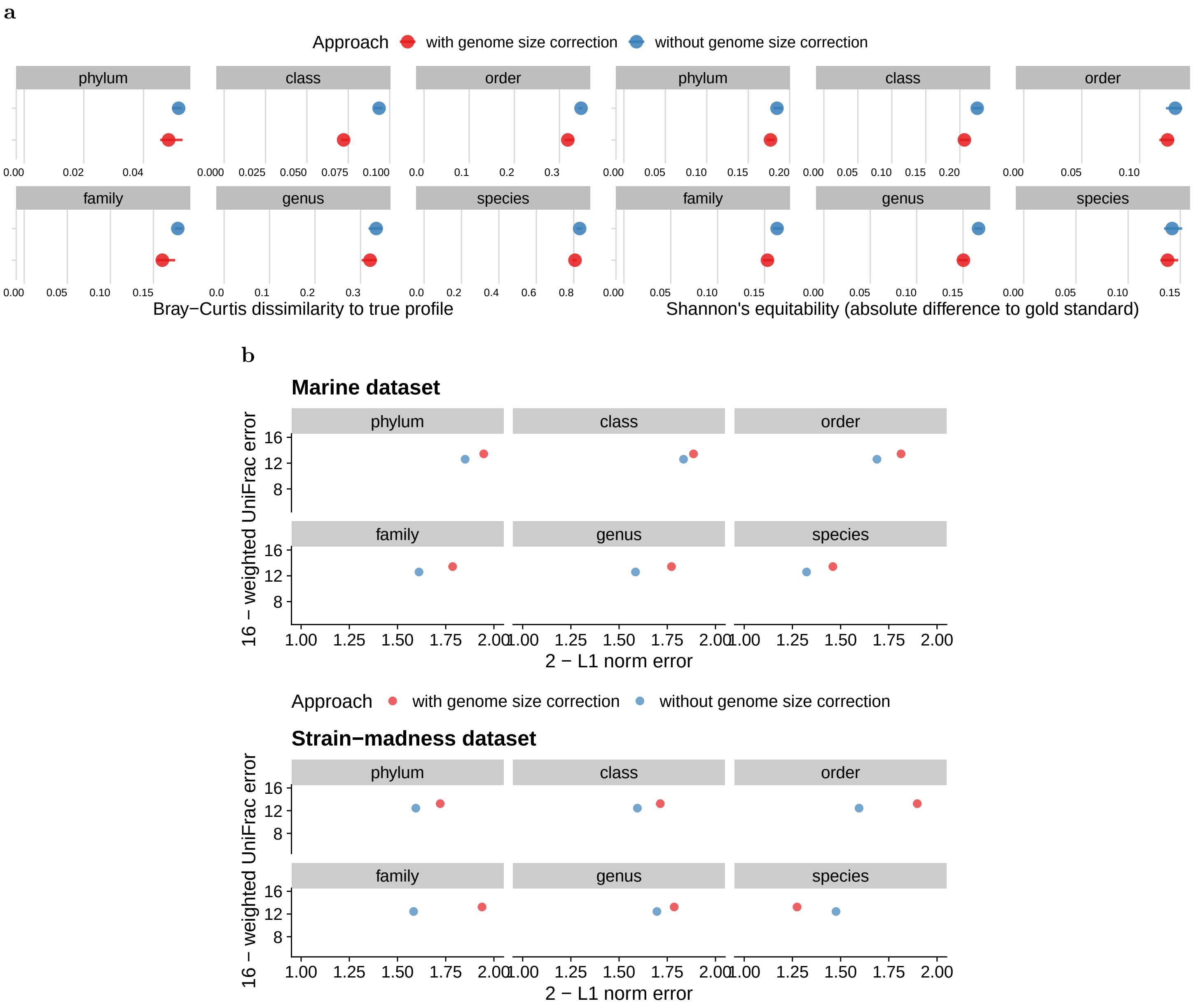


Figure S9: Effect of size correction on the abundance profiling performance. Profiles without size correction correspond to the vector $p_t^r$, and $\hat{p}_t^r$ is the size-corrected version. (a) Taking genome sizes into consideration results in lower Bray-Curtis dissimilarity to the true profile for all ranks in the CAMI-I high-complexity samples. (b) Similarly for the CAMI-II datasets, size-corrected abundance profiles perform better according to both L1 norm error (except at the species rank for strain-madness dataset) and weighted UniFrac error.

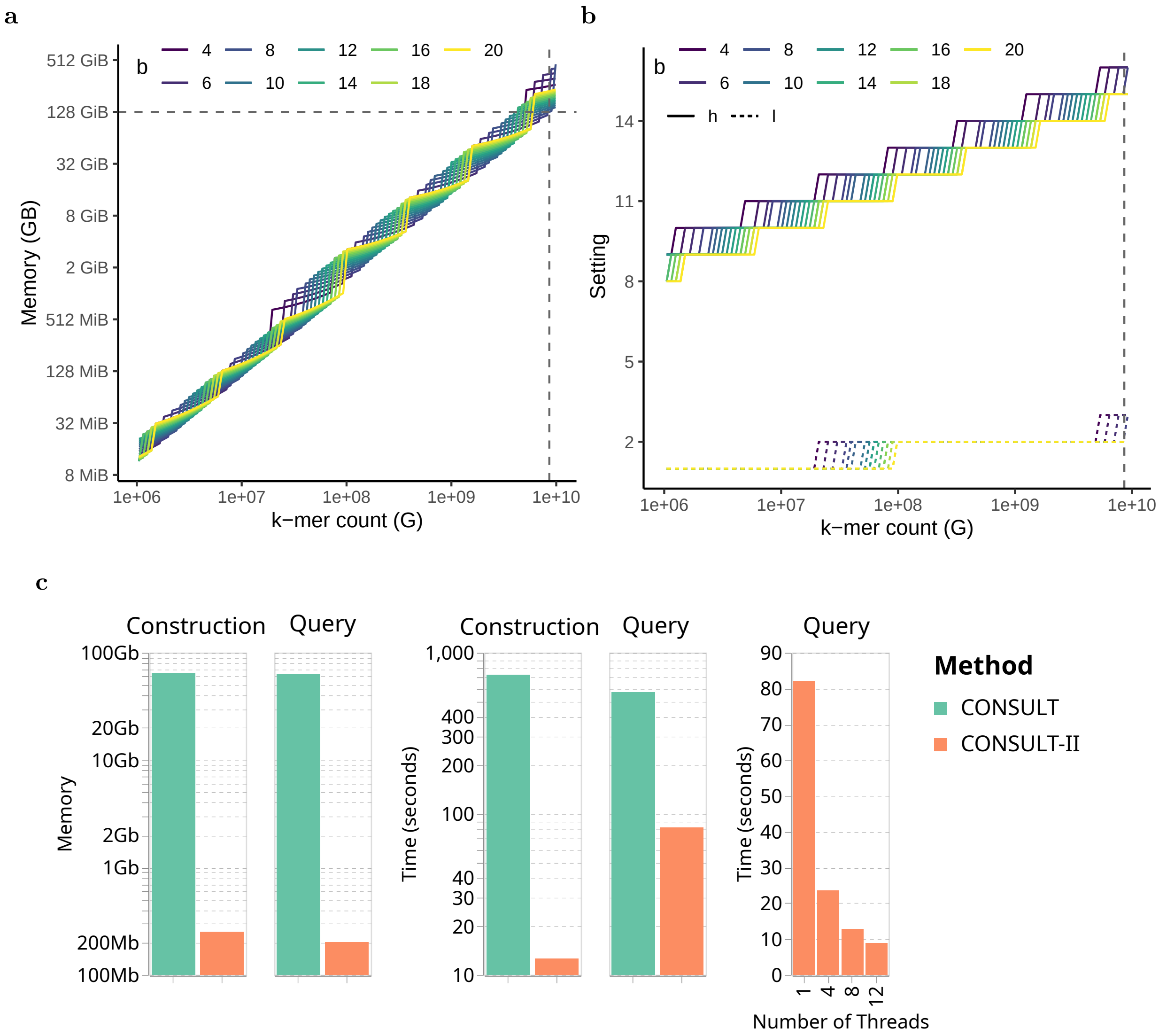


Figure S10: (b) Different configurations of $h$ and $l$ to accommodate $G$ $k$-mers ($x$-axis) and at the same time to achieve performance $\rho(3) = 0.45$ for various choices of $b$. (a) Fixing the performance according to part (b) results in different memory requirements for various $b$; note the log-scale y-axis, and hence optimizing $b$ can reduce memory requirements substantially. (c) Comparison of CONSULT and CONSULT-II in terms of computational resources. CONSULT-II achieves the same level of performance with CONSULT, but with much lower resource requirements, by using the heuristic described and setting $b$ to the optimal value. Reported memory usages and runtimes are measured in a real dataset, consisting of six E. coli genomes in the reference set and four E. coli genomes in the query set (12 million 32-mers and 1 million 150bp reads, respectively).

### References


[1] W. Huang, L. Li, J. R. Myers, and G. T. Marth. ART: A next-generation sequencing read simulator. *Bioinformatics*, 28(4):593–594, 2012. ISSN 13674803. doi: 10.1093/bioinformatics/btr708.

[2] H. Li. Seqtk, toolkit for processing sequences in FASTA/Q formats., 2018. URL `https://github.com/lh3/seqtk`.

[3] J. Lu, F. P. Breitwieser, P. Thielen, and S. L. Salzberg. Bracken: estimating species abundance in metagenomics data. *PeerJ Computer Science*, 3:e104, Jan. 2017. ISSN 2376-5992. doi: 10.7717/peerj-cs.104. URL `https://doi.org/10.7717/peerj-cs.104`.

[4] G. Marçais and C. Kingsford. A fast, lock-free approach for efficient parallel counting of occurrences of k-mers. *Bioinformatics*, 27(6):764–770, 2011. ISSN 1367-4803. doi: 10.1093/bioinformatics/btr011. URL `https://doi.org/10.1093/bioinformatics/btr011`.

[5] F. Meyer, A. Bremges, P. Belmann, S. Janssen, A. C. McHardy, and D. Koslicki. Assessing taxonomic metagenome profilers with OPAL. *Genome Biology*, 2019. ISSN 1474760X. doi: 10.1186/s13059-019-1646-y.

[6] F. Meyer, A. Fritz, Z.-L. Deng, D. Koslicki, T. R. Lesker, A. Gurevich, G. Robertson, M. Alser, D. Antipov, F. Beghini, D. Bertrand, J. J. Brito, C. T. Brown, J. Buchmann, A. Buluç, B. Chen, R. Chikhi, P. T. L. C. Clausen, A. Cristian, P. W. Dabrowski, A. E. Darling, R. Egan, E. Eskin, E. Georganas, E. Goltsman, M. A. Gray, L. H. Hansen, S. Hofmeyr, P. Huang, L. Irber, H. Jia, T. S. Jørgensen, S. D. Kieser, T. Klemetsen, A. Kola, M. Kolmogorov, A. Korobeynikov, J. Kwan, N. LaPierre, C. Lemaitre, C. Li, A. Limasset, F. Malcher-Miranda, S. Mangul, V. R. Marcelino, C. Marchet, P. Marijon, D. Meleshko, D. R. Mende, A. Milanese, N. Nagarajan, J. Nissen, S. Nurk, L. Oliker, L. Paoli, P. Peterlongo, V. C. Piro, J. S. Porter, S. Rasmussen, E. R. Rees, K. Reinert, B. Renard, E. M. Robertsen, G. L. Rosen, H.-J. Ruscheweyh, V. Sarwal, N. Segata, E. Seiler, L. Shi, F. Sun, S. Sunagawa, S. J. Sørensen, A. Thomas, C. Tong, M. Trajkovski, J. Tremblay, G. Uritskiy, R. Vicedomini, Z. Wang, Z. Wang, Z. Wang, A. Warren, N. P. Willassen, K. Yelick, R. You, G. Zeller, Z. Zhao, S. Zhu, J. Zhu, R. Garrido-Oter, P. Gastmeier, S. Hacquard, S. Häußler, A. Khaledi, F. Maechler, F. Mesny, S. Radutoiu, P. Schulze-Lefert, N. Smit, T. Strowig, A. Bremges, A. Sczyrba, and A. C. McHardy. Critical Assessment of Metagenome Interpretation: the second round of challenges. *Nature Methods*, 19(4):429–440, Apr. 2022. ISSN 1548-7091, 1548-7105. doi: 10.1038/s41592-022-01431-4. URL `https://www.nature.com/articles/s41592-022-01431-4`.

[7] B. D. Ondov, T. J. Treangen, P. Melsted, A. B. Mallonee, N. H. Bergman, S. Koren, and A. M. Phillippy. Mash: fast genome and metagenome distance estimation using MinHash. *Genome Biology*, 17(1):132, Dec. 2016. ISSN 1474-760X. doi: 10.1186/s13059-016-0997-x.

[8] R. Ounit, S. Wanamaker, T. J. Close, and S. Lonardi. CLARK: fast and accurate classification of metagenomic and genomic sequences using discriminative k-mers. *BMC Genomics*, 16(1):236, Dec. 2015. ISSN 1471-2164. doi: 10.1186/s12864-015-1419-2. URL `http://www.biomedcentral.com/1471-2164/16/236`.

[9] A. O. B. Şapcı, E. Rachtman, and S. Mirarab. Consult-ii: Taxonomic identification using locality sensitive hashing. In K. Jahn and T. Vinař, editors, *Comparative Genomics*, pages 196–214, Cham, 2023. Springer Nature Switzerland. ISBN 978-3-031-36911-7.

[10] D. E. Wood, J. Lu, and B. Langmead. Improved metagenomic analysis with Kraken 2. *Genome Biology*, 20(1):257, Dec. 2019. ISSN 1474-760X. doi: 10.1186/s13059-019-1891-0. URL `https://genomebiology.biomedcentral.com/articles/10.1186/s13059-019-1891-0`.
